# De novo protein design by inversion of the AlphaFold structure prediction network

**DOI:** 10.1101/2022.12.13.520346

**Authors:** Casper Goverde, Benedict Wolf, Hamed Khakzad, Stéphane Rosset, Bruno E. Correia

**Author notes:** These authors contributed equally (Casper Goverde, Benedict Wolf).

## Abstract

*De novo* protein design enhances our understanding of the principles that govern protein folding and interactions, and has the potential to revolutionize biotechnology through the engineering of novel protein functionalities. Despite recent progress in computational design strategies, *de novo* design of protein structures remains challenging, given the vast size of the sequence-structure space. AlphaFold2 (AF2), a state-of-the-art neural network architecture, achieved remarkable accuracy in predicting protein structures from amino acid sequences. This raises the question whether AF2 has learned the principles of protein folding sufficiently for de novo design. Here, we sought to answer this question by inverting the AF2 network, using the prediction weight set and a loss function to bias the generated sequences to adopt a target fold. Initial design trials resulted in de novo designs with an overrepresentation of hydrophobic residues on the protein surface compared to their natural protein family, requiring additional surface optimization. In silico validation of the designs showed protein structures with the correct fold, a hydrophilic surface and a densely packed hydrophobic core. In vitro validation showed that several designs were folded and stable in solution with high melting temperatures. In summary, our design workflow solely based on AF2 does not seem to fully capture basic principles of de novo protein design, as observed in the protein surface’s hydrophobic vs. hydrophilic patterning. However, with minimal post-design intervention, these pipelines generated viable sequences as assessed experimental characterization. Thus such pipelines show the potential to contribute to solving outstanding challenges in de novo protein design.

## Introduction

De novo protein design aims to create stable, well-folded proteins with sequences distant from those found in nature and potentially new functions. To date, de novo proteins remain challenging to design because the solution space expands exponentially with each additional amino acid in the sequence. Therefore, it is crucial to develop new computational methods that capture the underlying principles that govern protein structure, allowing for efficient exploration of the structure-sequence space and the design of more complex protein folds and functions.

To date, computational protein design has seen significant advances in creating proteins with novel folds and functionalities such as enzymes (Jiang et al., 2008; Röthlisberger et al., 2008), protein-protein interactions (Gainza et al., 2022; Koday et al., 2016; Marchand et al., 2022), protein switches (Giordano-Attianese et al., 2020; Langan et al., 2019) and vaccines (Correia et al., 2014; Sesterhenn et al., 2020; C. Yang et al., 2021).

A classical approach in computational design is fixed backbone design where a novel sequence is fitted to an existing protein topology from the Protein Data Bank (PDB) (Berman et al., 2000). Backbone and side-chain rotamer conformations of the residues are sampled and scored with various scoring functions allowing the creation of new protein structures and functions such as zinc finger domains (Dahiyat & Mayo, 1997), protein sensors (Feng et al., 2015; Glasgow et al., 2019), enzymes (Jiang et al., 2008; Röthlisberger et al., 2008; Siegel et al., 2010) and small molecule binders (Tinberg et al., 2013).

Fragment assembly design methods have been extensively used to generate diverse backbones from scratch. This method assembles structural protein fragments into the desired fold which has proven successful in the design of de novo beta barrels (Dou et al., 2018), TIM barrels (Huang et al., 2016), jellyroll structures (Marcos et al., 2018) and various alpha-beta proteins (Correia et al., 2014; Koga et al., 2012). Fragment assembly has also successfully designed novel protein folds that were not found in nature (Kuhlman et al., 2003).

De novo proteins generated with fragment assembly have also been functionalized by constructing topologies that stabilize a functional motif. This method has resulted in the design of biosensors and vaccine candidates (Sesterhenn et al., 2020; C. Yang et al., 2021). However, this method is completely dependent on fragments extracted from native proteins that are included in limited libraries and are also notoriously inefficient in generating good quality backbones if structural constraints are not applied.

Recently, de novo protein design has taken an exciting turn with the emergence of deep learning tools for protein modeling, allowing the generation of proteins without relying on fragment libraries that explore diverse solutions of the sequence-structure space. With an increase in sequence and structural data combined with significant progress in deep learning, these techniques have transformed protein structure prediction and design (Ovchinnikov & Huang, 2021). For example, deep learning has proven to be an excellent tool for sampling the backbone conformational space through, e.g., Generative Adversarial Networks (GANs) (Anand et al., 2022) and Variational Auto-Encoders (VAEs) (Eguchi et al., 2022; Guo et al., 2021). Deep learning methods can also be used for sequence generation given a target backbone; through the interpolation between sequence and structural data, sequences that fit a target topology can be found (Anand et al., 2022; Dauparas et al., 2022; Ingraham et al., 2019).

The trRosetta structure prediction network (J. Yang et al., 2020) was recently applied for sequence generation given a target structure. This was achieved by using the error gradient between the sequence and the target contact map to optimize a Position Specific Scoring Matrix (PSSM) (Henikoff & Henikoff, 1996; Norn et al., 2020). Protein folding network sequence generation was followed by removal of the structure constraints and only optimizing for protein stability. This method allows the network to ‘hallucinate’ a sequence and structure different from those in nature (Anishchenko et al., 2021). Hallucination involves optimizing a randomly initialized amino acid sequence to a loss function. Experimental data confirmed that several hallucinated proteins adopted the target fold. Additionally, it was shown that a scaffold could be hallucinated to stabilize a functional site, supporting the desired structure (Tischer et al., 2020). The latest version of this hallucination pipeline uses a more accurate prediction pipeline using RoseTTAfold (RF) (Baek et al., 2021). Additionally, an inpainting step was used to optimize designs by randomly masking and predicting the most likely amino acids. This method successfully designed de novo proteins with active sites, epitopes and protein binding sites (Wang et al., 2022).

Recently, AlphaFold2 (AF2), an end-to-end structure prediction network, reached unprecedented accuracy levels close to experimental methods for structure determination (Jumper et al., 2021). AF2 predicts 3D atom coordinates of the protein structure given a Multiple Sequence Alignment (MSA) and structures of homologs (templates). Although the network was trained on MSAs containing co-evolutionary signals and templates as inputs, AF2 can accurately predict protein structures of de novo proteins from a single sequence alone (Pereira et al., 2021). This indicates that AF2 can generalize to de novo designed protein sequences, potentially providing a new tool for protein design.

New ideas for harnessing the power of AF2 followed shortly after its release. The first attempts started with an initialized sequence from a generative model and applied a greedy algorithm to optimize the designs for various loss functions (Jendrusch et al., 2021; Moffat et al., 2021). These AF2-based design pipelines were used to design monomers, oligomers and protein switches which were validated using various in silico metrics. In recent research, it was also demonstrated how symmetric protein homooligomers could be designed through the help of deep network hallucination using AF2. A backbone was hallucinated using AF2 and the sequence is redesigned using protein MPNN, a message-passing graph neural network that improves design success rates (Dauparas et al., 2022; Ingraham et al., 2019; Wicky et al., 2022). However, the conformational search of these pipelines is solely based on stochastic MCMC methods and is, therefore, computationally expensive.

We hypothesized that we could devise an efficient design strategy by inverting the AF2 structure prediction network. Hence, a structural loss is backpropagated through AF2 to generate amino acid sequences compatible with a target fold. Through error gradient backpropagation combined with MCMC optimization, we explored several protocols to generate sequences given a target structure using AF2-based pipelines we call AF2-design. We analyzed the generated sequences both *in silico* and *in vitro*, showing that our AF2-based protocol can be leveraged for de novo protein design.

## Results

### AF2-design methodological approach

The general goal of protein structure prediction networks such as AF2 is to predict the tertiary structure given the sequence. Here, we propose using AF2 for the inverse problem, generating protein sequences given a target backbone. One inversion strategy is error backpropagation (Simonyan et al., 2014), where given a target protein structure, the input sequence is optimized to a loss function. Inspired by the trDesign method (Norn et al., 2020), we developed an AF2-design pipeline as illustrated in **Figure 1a**. The first step was to initialize an input sequence and predict its structure. Since we started with non-natural sequences, natural homologs are unavailable and, therefore, MSAs and structural templates are disabled in the AF2 network (also referred to as single sequence mode).

**Figure 1:**
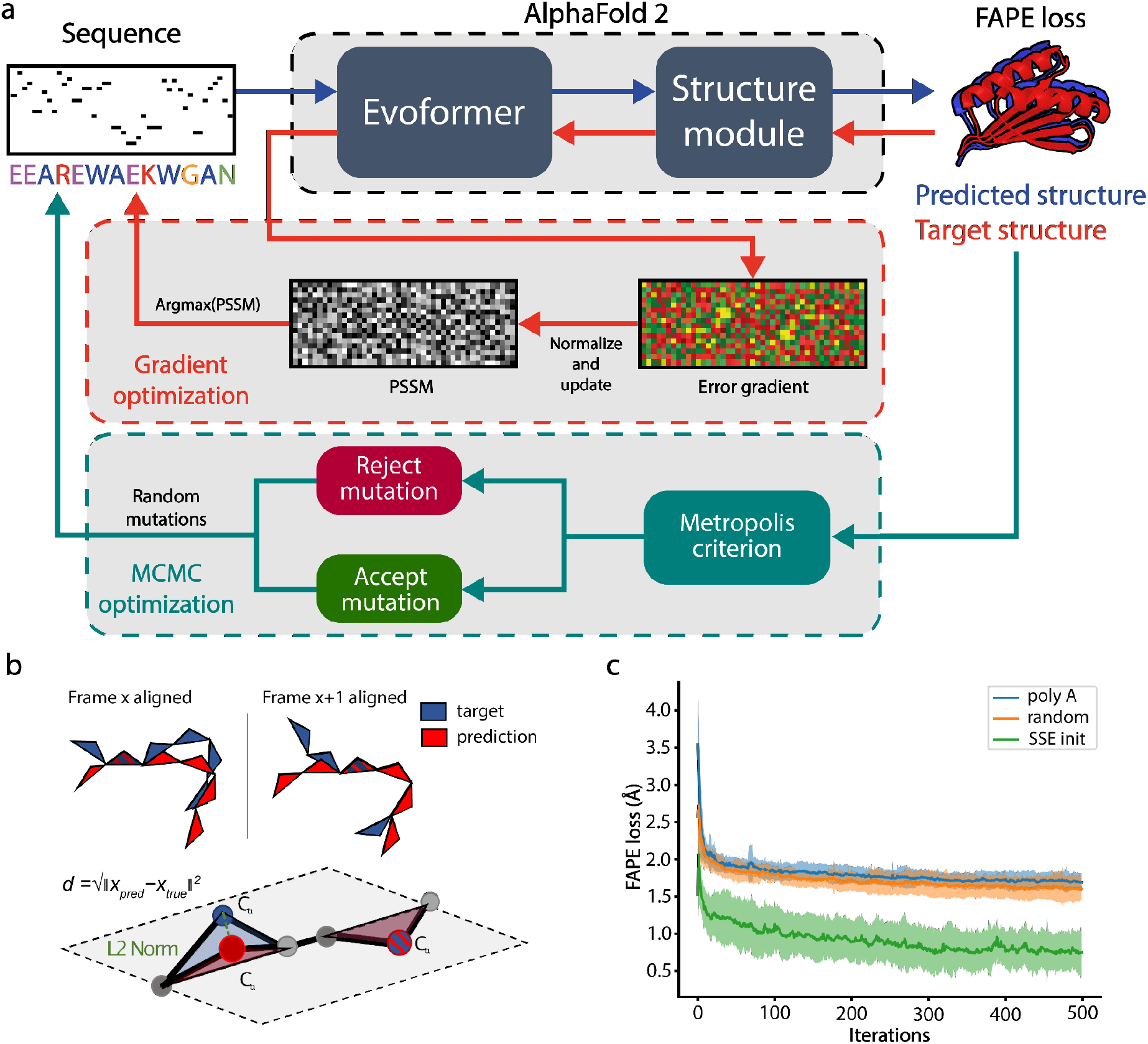
AlphaFold-based pipeline for sequence design. **a)** The design pipeline consists of two sequence optimization methods, GD and MCMC-based optimization. In the GD optimization step, the design process is initialized with a secondary-structure biased sequence, where the structure is predicted upon each sequence change (blue arrows). Next, sequence information from the AF2 network is extracted by backpropagating the error gradient between the predicted and desired target structures (red arrows). This error gradient is then used to optimize a predicted PSSM where the amino acids with the highest probability are used as the input to the next optimization round. After several rounds of gradient-based optimization, the sequence with the lowest error (FAPE loss) is selected and used for further MCMC-based optimization to decrease the distance between the target and the predicted protein structure (green arrows). A random mutation is introduced to the sequence in the refinement step and accepted based on the Metropolis criterion. **b)** The FAPE loss is computed by taking the L2 norm (Euclidean distance) between the C-alpha coordinates after alignment to each of the residue frames.. **c)** Examples of design trajectories using different initialization sequences. SSE sequence initialization was the only approach that converged to the correct folds within 500 iterations.

After predicting the structures of these sequences, the structural error is computed using the Frame Aligned Point Error (FAPE) (Jumper et al., 2021). The FAPE loss is a rotationally invariant form of the root mean square deviation (RMSD). The backbone atoms (frames) are aligned and the interatomic C-alpha distances between the predicted and target structure are computed (**Figure 1b**). Since the FAPE loss was the primary loss component during training, we reasoned that backpropagating this loss would result in the most accurate generation of protein sequences.

AF2 consists of five networks trained on different parameters to generate five structural models. We adapted the original AF2 pipeline so all networks run in parallel on a single GPU, increasing prediction speed fivefold. Additionally, using all AF2 models prevents overfitting to a single model and thus designs a sequence all models agree upon. Next, the error gradients of the one-hot-encoded input sequence are computed and averaged to obtain an Nx20 mean error gradient matrix where N is the sequence length. This error gradient measures the contribution of each residue to the loss. Using a gradient descent (GD) algorithm we use the error gradient to update a PSSM, representing the probability distribution of the amino acids for each position. After each round, the most probable amino acids are selected from the PSSM and used in the next iteration as the input. To test the performance of AF2-design, we sought to de novo design a collection of protein folds. Specifically, we attempted to generate new sequences that would adopt the folds of top7, protein A, protein G, ubiquitin and a 4-helix bundle (**Figure S1 and Table S1**). Throughout our design simulations we observed that the initial input sequence significantly impacts the convergence rate of AF2-design. When initializing the GD pipeline with a poly-alanine or a random sequence, the network did not converge within 500 iterations in all folds (**Figure 1c**). One potential reason is that the predicted structure of a poly-alanine is disordered and thus results in a very noisy error gradient with no smooth trajectory to converge into a folded conformation. We hypothesized that this problem could be solved by introducing sequences that favor the local secondary structure propensity; hence we proposed a Secondary Structural Element (SSE) initialization strategy (**Figure S2**). In the starting sequence, the amino acid identities are assigned according to which SSE they are expected to form: helical residues get assigned alanines, betasheets valines and loops with glycines. This strategy led to convergence to the correct fold within 500 iterations for all the tested design targets. However, since AF2 is deterministic in inference mode, the same designed sequence would be obtained for each run. To address this issue, we mutate 10% of the amino acids in each design simulation in the starting sequence, resulting in the generation of diverse sequences. Additionally, we masked cysteines in the PSSM, excluding these amino acids from sequence design to avoid an overrepresentation of cysteine residues and disulfide bonds.

In addition to GD optimization, we also implemented a Markov Chain Monte Carlo (MCMC) search algorithm. We observed that GD allows for a quick convergence within the sequence space, but further improvement in TM scores could be achieved (Zhang & Skolnick, 2004, 2005) by adding an MCMC optimization step (**Figure 1a and 2a**). We hypothesized that this extra MCMC step allows the escape from local minima where GD converges. During MCMC optimization, four random positions in the sequence are chosen and mutated every iteration. We sample amino acids from a probability distribution of natural protein compositions for the mutations, identical to the trDesign approach (Anishchenko et al., 2021). Since the computationally expensive error backpropagation step is omitted in this step, we can increase the number of passes (recycles) through the network without making the runtime unfeasible, resulting in an increased structure prediction quality. Finally, the designs with the lowest FAPE loss are selected and relaxed in an AMBER force field to remove interatomic clashes (Hornak et al., 2006).

**Figure 2:**
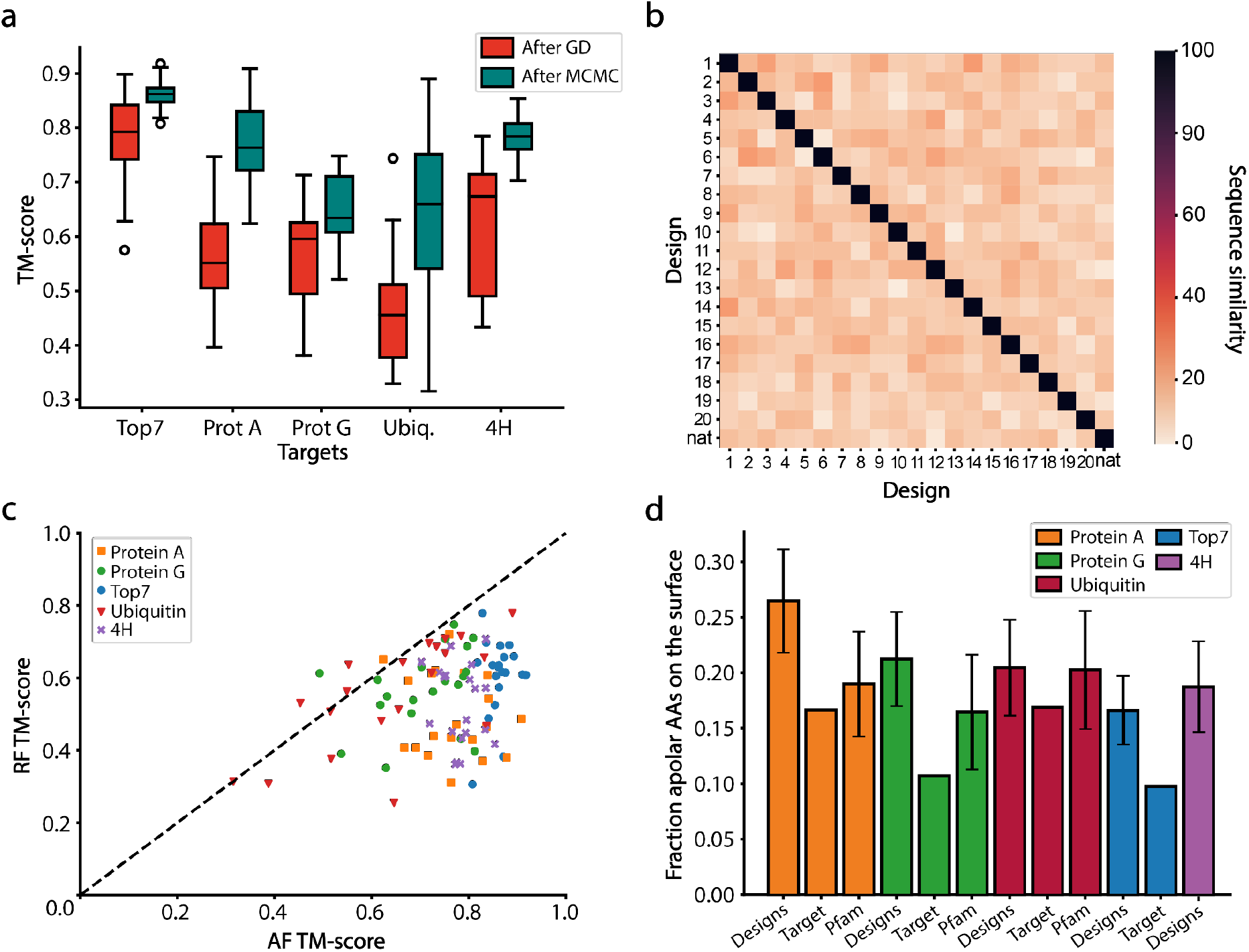
Features of AF2 designed sequences. a) TM-scores of designed sequences using a combination of GD and MCMC optimization. Red - TM-scores after GD design; Green - TM-scores after an additional round of MCMC sampling. For each fold, 20 rounds of GD and MCMC optimization were performed. MCMC optimization significantly improves the TM scores of the GD based designs. b) Evaluation of the sequence diversity obtained within the top7 designs. The designed sequences have a low sequence similarity (between 10-30%) when compared to one another and to the native sequence. c) Structure prediction of the AF2 designed sequences using AF2 and RF. In most instances, RF predicts lower TM-scores than AF2. d) Fraction of hydrophobics on the surface before the surface redesign step. All the designs have more hydrophobic residues on their surface than their target fold. When comparing the designs to their protein family, we find that the designs of protein A and protein G have slightly more hydrophobics on their surface than similar folds found in nature. Top7 and 4H are de novo proteins hence do not have a protein family, additionally, 4H is a backbone model designed with the TopoBuilder and as such there is no sequence to be compared to.

An additional surface design step was performed with Rosetta to increase solubility and prevent aggregation by removing solvent-exposed hydrophobic amino acids. Accordingly, surface residues are randomly mutated to hydrophilic amino acids and optimized using the Rosetta energy function (Alford et al., 2017). The refined sequences are then predicted using both AF2 and RF. The final designs are selected based on both TM-score (> 0.6 for AF2 and 0.5 for RF-generated models) and AF2 confidence in the predicted structure (AF2 plddt > 60).

### Computational evaluation of designed sequences

To test the performance of AF2-design, five distinct folds were selected as design targets. The attempted folds were: top7, one of the first de novo proteins (Kuhlman et al., 2003); protein A, ubiquitin, and protein G, small globular folds; 4H, a 4-helix bundle designed by TopoBuilder (C. Yang et al., 2021) (**Figure S1**). For each target fold, 20 designs were generated using 500 iteration trajectories of GD and the fold similarity between designs and target is evaluated using TM-score. The sequences generated by GD largely present the correct fold (**Figure 2a**, red boxes).

Following the GD design stage the sequences were refined using MCMC optimization, which improved the TM-scores for all designs (**Figure 2a**, green boxes). However, most of the designs of protein G and ubiquitin still had relatively low TM scores (< 0.7). Overall the observed trend was that a pipeline composed of a combination of GD and MCMC stages yields designs closer to the target folds. To assess the sequence diversity of the generated designs, we performed an all against all comparison of the designed and native sequences and observed a low sequence similarity between 20-30% (**Figure 2b** and **Figure S3**). Additionally, we sought to evaluate the designed sequences for structural accuracy using structure prediction tools. We used RF, a structure prediction network developed and trained independently from AF2 (Baek et al., 2021) and compared the results to AF2 predictions. Generally, RF predictions of the designed sequences showed lower TM scores than AF2 (**Figure 2c**), this is likely related to the fact that the sequences were generated with AF2. As such, orthogonal prediction tools may be valuable help to identify the best designs. In further analysis of the designs, we noticed a slight overrepresentation of hydrophobic amino acids on the surface in 3 out of the 4 folds where comparisons could be established (**Figure 2d**). In general the presence of hydrophobic aminoacids at the surface of the protein is seen as unfavorable for the solubility of the designs. Hence, a Rosetta surface redesign step was performed on the AF2-designed sequences to correct the exposed surface hydrophobics. The resulting sequences were validated by AF2 and RF predictions and the best designs were selected for experimental characterization. Comparing the selected models with the pre-surface redesign models we observe an increase in BLAST e-values (**Figure. S4**).

The designs characterized experimentally were generated both by GD-MCMC optimization as well as MCMC-only design pipelines, trajectories are shown in **Figures S5 and S6**. Next, the designs were filtered on AF2 TM scores > 0.6, RF TM scores > 0.5 and AF2 confidence (pLDDT) > 60 of the predicted structures resulting in 39 designs (**Table S1-3**). The predicted structures of the designs closely resemble their target folds as shown by structural superpositions **(Figures 3a, S7, S8)** and the TM scores (**Figure 3b**). At the sequence level, the designs explore novel sequence space as observed by the high E-values derived from sequence alignments performed with BLAST on non-redundant protein sequences (**Figure 3c**) (Gish & States, 1993). In summary, our AF2-design pipelines can efficiently explore the non-natural sequence space given a pre-defined fold and can be validated using orthogonal structure prediction networks.

**Figure 3:**
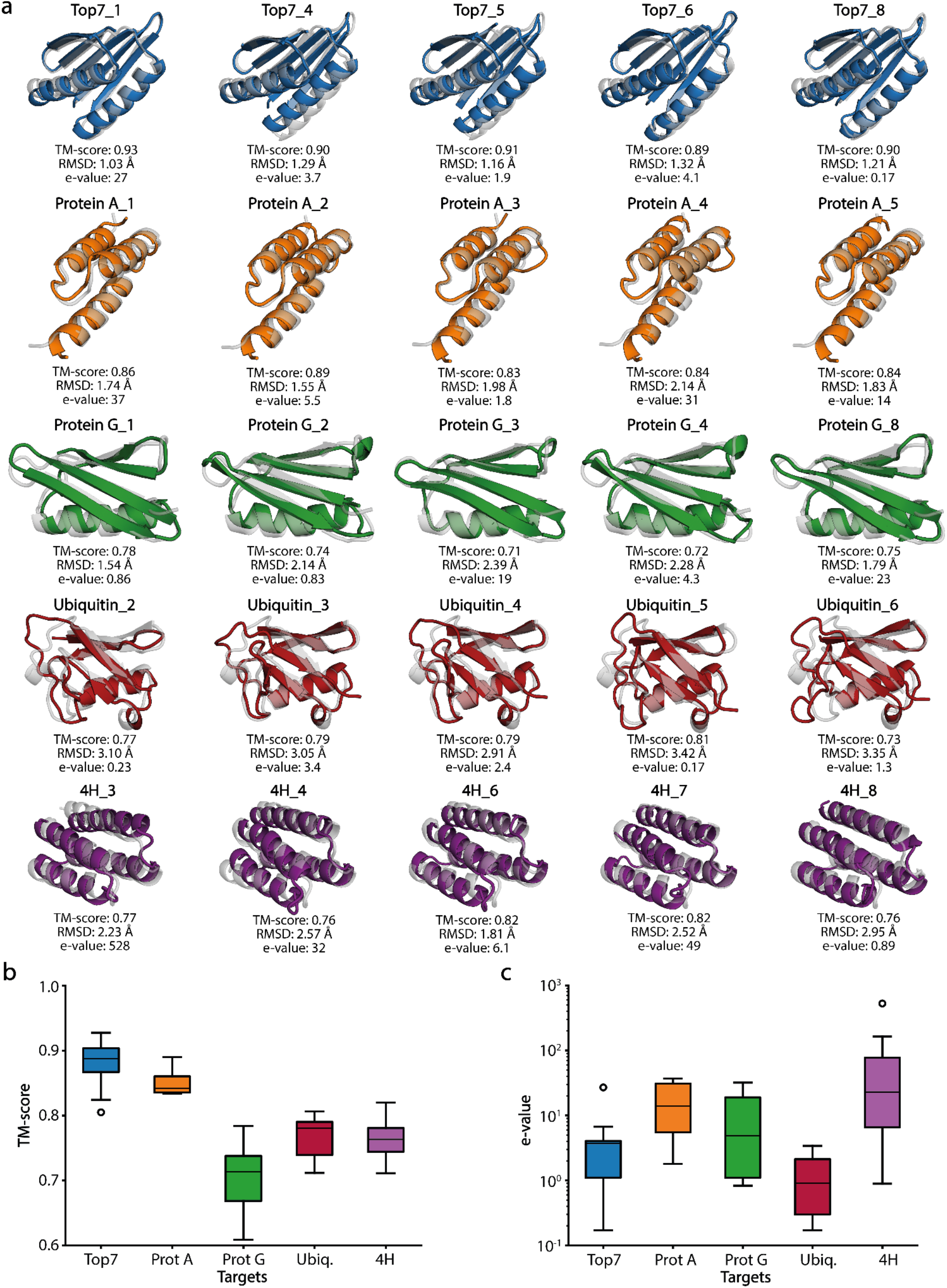
Overview of the structural and sequence properties of the AF2 generated designs. a) Alignment of the best design prediction vs. the reference structure (gray) according to the TM-score. top7 - blue, Protein A - orange, Protein G - green, Ubiquitin - red and 4H - purple. b) TM-scores of the final designs with a minimum above 0.60. c) The distribution of the nearest e-value of all designs.

### *In vitro* characterization of designed sequences

We next sought to biochemically characterize the AF2-designed sequences. Thirty-nine designs were cloned and expressed in E. coli (11 top7, 5 protein A, 9 protein G, 6 ubiquitin and 8 4H designs). Twenty-five designs were expressed solubly and purified by affinity and size exclusion chromatography (SEC) (3 top7, 5 protein A, 7 protein G, 6 ubiquitin and 4 4H designs). Ultimately only 3 of the 5 target folds yielded proteins with acceptable biochemical behavior (protein G, top7 and 4H). Several designs from these folds displayed monodisperse peaks indicating monomeric or dimeric states in solution as shown by SEC coupled to multi-angle light scattering (**Figure 4 and S9**). The oligomeric states observed in solution were single-species monomeric (4H_1) and mixed species between monomer/dimer (top7_1, protein G_1). Several designs showed similar secondary structure content to the original proteins, as assessed by CD spectroscopy. The designs based on de novo backbones (top7 and 4H) showed very high thermal stability with melting temperatures (T_m_) over 90 °C (**Figure 4**). The original top7 was also a hyper-stable protein (T_m_ > 90°C) (Kuhlman et al., 2003); this is an interesting observation given that the designs have only 12.4% of shared sequence identity, suggesting that such backbone may be prone to sequences conferring high thermal stability. The protein G designs showed cooperative unfolding with T_m_s of 49°C and 56 °C, respectively, which are considerably lower than the wildtype protein G (T_m_ = 89 °C) (Ross et al., 2001).

**Figure 4:**
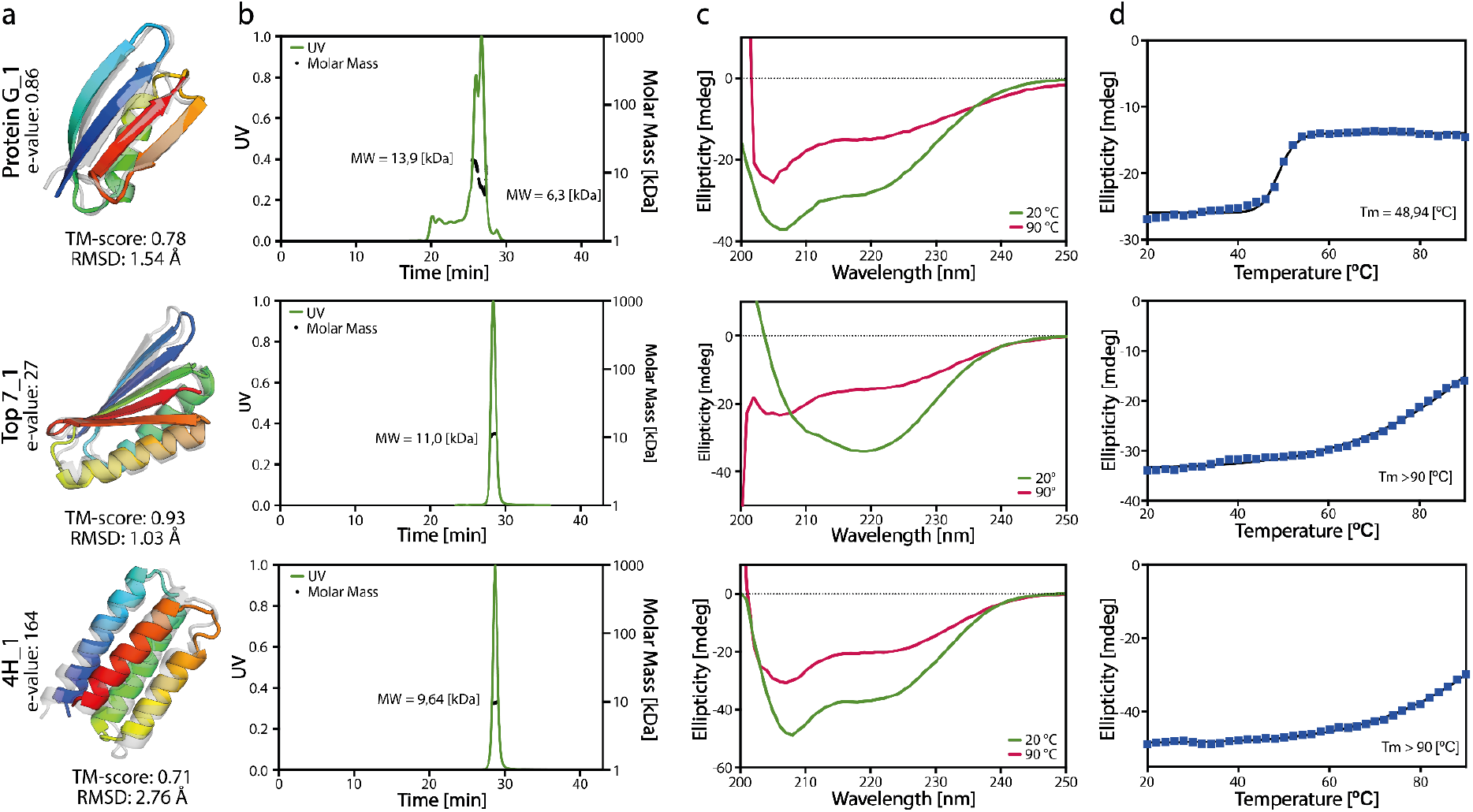
Experimental characterization of the designs. a) The superimposed structures of the design predicted by AF2 (colored) vs. the original (gray) structure. b) The SEC-MALS measurements indicated that protein G_1 and top7_10 appear in monomeric and dimeric forms, whereas 4H_1 appear purely as a monomer. The expected Molecular Weight (MW) of the designs was 7.7 kDa for protein G_1, 11.7 kDa for protein top7_10 and 11.2 kDa for protein 4H_1. c) The CD spectra of the designs showed that the designs were folded. d) The temperature melting curves per design. All experimental characterizations of the folded designs can be found in **Figure S9**.

In light of the experimental results obtained, we sought to learn which metrics could be the best predictors for folded designs. We trained a simple logistic regression model for folded and non-folded classification based on TM-score, pLDDT, Rosetta energy score (ddG) and packing score. It is important to highlight that our dataset is extremely small and this analysis should be taken as merely indicative. We tested our model using stratified k-fold cross-validation in which the test set of each fold contained one positive and one negative example (for each fold, the test set contained a unique positive sample and a random negative sample). After training and validating our model, we obtained a mean training AUC of 0.86 (± 0.02) on the training set and 0.88 (± 0.33) on the testing set. Analyzing the logistic regression coefficients, we concluded that the most important features are AF2 and RF TM-score and AF2 pLDDT (**Figure 5a-c**). The RF TM-score showed the most significant impact on the folded classification. Unexpectedly, a high AF2 TM-score showed to correlate with the unfolded designs. This correlation most likely is an effect of filtering on AF2 TM-score in combination with our small sample size. These results indicate that increasing the RF TM-scores and AF2 pLDDT filters might increase success rates in obtaining folded designs in vitro. However, and as mentioned above, the number of designs is small and more computational and experimental data is necessary for more robust conclusions. When inspecting the per residue pLDDT of protein G and ubiquitin-like fold designs, we found regions of the designs with low confidence (ubiquitin) or noticeable lower on the unfolded designs compared to the folded ones (protein G, **Figure 5d**). This observation is consistent with the findings reported by Listov et al., where per residue pLDDT scores were also utilized as an indicator of problematic areas in designed proteins (Listov et al., 2022). In conclusion, we obtained eight soluble and well-folded proteins out of 39 designs. Our analysis points to some quality indicators derived from structure prediction networks for selecting folded designed sequences.

**Figure 5:**
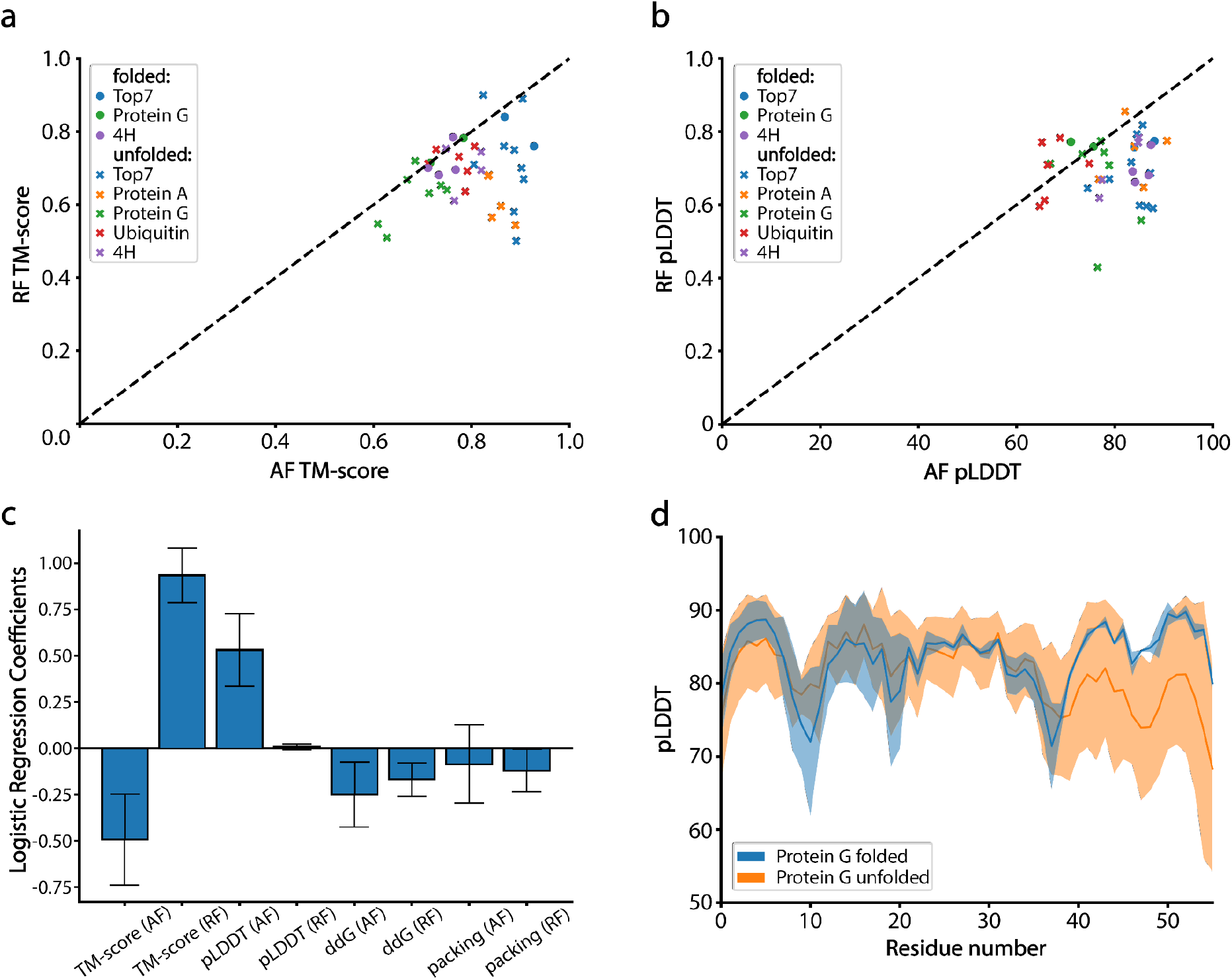
In silico analysis of folded and non-folded designs. a) TM-scores of AF2 predicted models vs. RF predicted models. b) Confidence in the models as predicted by AF2 and RF. c) Logistic regression coefficients for a model trained on features of all folded and non-folded designs. The AF2 and RF TM-score, and AF2 pLDDT have the strongest impact on the classification. d) Per residue confidence of protein G designs as predicted by AF2. For two of the non-folded designs, AF2 predicted the two beta sheets at the C-terminal end with lower confidence than the other beta strands in the structure. The per residue pLDDT for all designs can be found in **Figure S10-11**.

## Conclusion

Machine learning approaches have had a transformative impact on protein structure prediction. Historically, predicting structure from sequence has been considered an essential requirement for protein design endeavors. However, design exercises pose a much more complex challenge due to the vast sequence space and the restricted subset that encodes particular folds. With this challenge in mind, we tested if the state-of-the-art structure prediction network AF2 could be repurposed for design using several sequence-structure generation and selection workflows. Several important observations arose from our studies for the generated designs: I) the networks converged relatively fast (small number of steps) into the target folds and showed dependency on the starting sequence; II) a substantial amount of designs generated lacked the observed/expected hydrophobicity patterns in soluble proteins and displayed a high proportion of hydrophobic residues in solvent-exposed areas, despite this lack of agreement with the fundamental physicochemical properties of proteins the prediction network still produced structures very similar to those of the target folds; III) upon refinement of the sequences with Rosetta, specifically of surface positions, several designs are biochemically well behaved, folded and stable in solution. Our results demonstrate that a combination of AF2 and RosettaDesign can be used for de novo protein design. In contrast to previously developed AF2 design methods, we present an approach that utilizes error gradients to achieve quick convergence. With this protocol, we show that it is possible to explore non-natural sequence spaces given a protein topology which is useful for the design of new functional proteins. Specifically, such functional design can be achieved within our pipeline by adding sidechain losses allowing to control the configuration of binding sites, as has also been shown by Wang et al. (Wang et al., 2022). This framework has potential uses for designing small molecule binding sites, catalytic sites and protein-protein interactions (PPIs). Additionally, for PPI design, the ability of AF2 to predict multimers can be used to sample binder sequences that can be optimized given a positional loss and confidence through error backpropagation. As shown by Wicky et al., AF2-based design pipelines can also be used for sampling the unexplored protein structural space. Using this same loss function, we expect to be capable of doing the same but considerably faster since we make use of error backpropagation. As a next step, we would like to add several additional components to the loss function such as plddt and ptm scores. This should increase the success rate of our designs and allow the fast design of complex protein folds. To conclude, this work demonstrates that inversion of structure prediction networks allows for the design of de novo proteins and hold promise for tackling complex problems such as protein binder and functional site design.

## Methods

### Loss function

AF2’s loss function was built of several components of which the FAPE loss was the major contributor for the training of the structural module. The FAPE loss measures the L2 norm (Euclidean distance) of all C-alpha atoms compared to the ground truth under many different alignments, making the loss invariant to rotations and translations. Hence, we chose the FAPE loss to calculate the error gradients guiding our sequence search (Jumper et al., 2021). The FAPE loss can be regularized with a clamped L1 norm, restricting the FAPE loss to a maximum value of 10 Å between two atom positions. This results in a loss more focused on accurate local positioning and de-emphasizing longer range errors.

### Gradient descent optimization

We initialized the target amino acid sequences based on the secondary structure of the residue derived from the target fold. The secondary structure assignments were then encoded in sequences using alanines for helix, valines for beta sheet and glycines for loop residues. This introduces a bias towards the correct local structure, aiding faster convergence of the design trajectories. To ensure sequence diversity of the generated designs we randomly mutate 10% of the amino acids in the initial sequence of each design trajectory.

The starting sequence is then one-hot-encoded (OHE) and passed through the AF2 network without any recycles. AF2 consists of five differently tuned networks to generate five structural models, in our design pipeline we make use of all the five generated models. Next, we compute the mean Frame Aligned Point Error (FAPE) of the models and calculate the mean error gradient of the OHE input. An empty Position Specific Scoring Matrix (PSSM) of shape Nx20 is initialized and a value of 0.01 is assigned to the input residues. We used the ADAM optimizer (Kingma & Ba, 2017) to update the PSSM with the normalized gradient which, after softmaxing the matrix, results in a probability distribution of the amino acids per position. Next, the most probable amino acid identities per position are determined and used to generate the new input sequence for the next iteration. Additionally, we set the values for cysteine in the PSSM to negative infinity resulting in designed sequences without cysteines.

### Markov Chain Monte Carlo optimization

As starting sequences we use the designs from GD runs or SSE initialized sequences. In each iteration, we mutate four random residues in the sequence to an amino acid sampled from a natural amino acid distribution (Anishchenko et al., 2021). We predict the structure using AF2 with 3 recycles, enabling more accurate predictions (Mirdita et al., 2022). Next, we calculate the mean FAPE loss between the predicted structures and target fold and use a Metropolis-Hastings algorithm to accept or reject the mutated sequence (Hastings, 1970). If the introduced mutations result in a reduction of the FAPE loss, the new sequence is accepted. If not, the sequence can be accepted based on the following metropolis criterion:

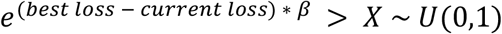

Where *X* is sampled from a uniform distribution between 0 and 1 and *β* is set to an initial value of 80 and doubled every 250 steps, making it unlikely that structures with a much higher loss are accepted. This Metropolis-Hastings search allows us to escape from local minima during our exploration of the sequence space.

### Hardware settings

For the design pipeline one Nvidia Tesla V100 (32GB) was used. The prediction of a protein with a sequence length of 92 (top7) and calculation of the gradients takes ~6 seconds for one iteration without recycling. For MCMC optimization one iteration took ~8.5 seconds using 3 recycles.

### Model settings

We initialized our AF2 network configurations using the settings of ‘model_5_ptm’. Additionally, since we run AF2 in single sequence mode, we disabled MSAs and template processing in the settings, reducing runtime and memory usage. The network runs in deterministic mode to ensure reproducible results, meaning that dropout layers are disabled. Finally, we run all five components of the AF2 network in parallel, speeding up the prediction time fivefold in comparison to the original AF2 pipeline.

### Computational design protocol

The five protein folds chosen as design targets were: top7 (1QYS, 2.5 Å X-ray) - a 92 residue de novo protein with a fold unknown to nature; protein A (1DEE, 2.7 Å X-ray) - a small three helix bundle with 54 residues; protein G (1FCL, NMR) - a mixed alpha/beta protein with 56 residues; ubiquitin (1UBQ, 1.8 Å X-ray) - a mainly beta protein with 76 residues; 4H (Rosetta model) - a de novo design with 84 residues constructed using TopoBuilder (C. Yang et al., 2021) and Rosetta FunFolDes (Bonet et al., 2018).

We employed our AF2-design pipeline using 500 rounds of GD and 1000-6000 steps of MCMC optimization. Of the experimentally validated and folded designs 1QYS_1 and 1QYS_10 were obtained purely through MCMC optimization.

After AF2 sequence generation a Rosetta fixed backbone design protocol was used to redesign the surface residues. The surface residues were defined as residues with Solvent Accessible Surface Area (SASA) > 40 Å^2^, and were allowed mutate to hydrophilic or charged amino acids common in their respective secondary structural element: “DEHKPQR” for helices, “EKNQRST” for beta sheets and “DEGHKNPQRST” for loops. For relaxation and scoring of the designs the REF15 energy function was used (Alford et al., 2017). The TM-scores and C-alpha RMSDs between design and target structure were determined using TM-align (Zhang & Skolnick, 2005). The sequence similarity is defined as the number of positions at which the corresponding residue is different. The validation using roseTTAfold was performed without MSAs or structural templates.

### Protein expression and purification

The designs were ordered as synthetic gene fragments from Twist Bioscience with the addition of a C-terminal 6-His-Tag and cloned into a pET11b vector using NdeI and BlpI restriction sites. The designs were transformed into XL-10-Gold cells and the DNA was extracted and validated by Sanger sequencing. The validated DNA sequences were transformed into BL21 DE3 cells and put in 20 ml of LB medium with 100 μg/ml Ampicillin overnight at 37 °C as starting cultures. The next day, 500 ml of Auto-Induction medium with 100 μg/ml Ampicillin was inoculated with 10 ml overnight culture and grown at 37 °C to OD of 0.6 then the cultures were grown for ~20 h at 20 °C. Bacteria were pelleted by centrifugation and resuspended in lysis buffer (100 mM Tris-Cl pH 7.5, 500 mM NaCl, 5% Glycerol, 1 mM Phenylmethanesulfonyl fluoride, 1 μg/ml lysozyme and 1:20 of CelLyticT_m_ B Cell Lysis Reagent). The resuspensions were put at room temperature on a shaker at 40 rpm for 2 hours and then centrifuged at 48’300 g for 20 minutes. We filtered the supernatant with a 0.2 μm filter and loaded the mixture on a 5 ml HisTrapT_m_ FF column using an AKTApure system and a predefined method regarding Cytiva’s recommendations with that column. We used 50 mM Tris-HCl pH 7.5, 500 mM NaCl, 10 mM Imidazole as was buffer and processed the elution with 50 mM Tris-HCl pH 7.5, 500 mM NaCl, 500 mM Imidazole. We collected the main fraction released through the elution step and injected it on a Gel Filtration column Superdex 16/600 75 pg filled with PBS. The peaks corresponding to the size of the design were collected and concentrated to a concentration of approximately 1 mg/ml for further analysis. Folding and secondary structure content was assessed using circular dichroism in a Chirascan V100 instrument from Applied Photophysics. Melting temperature determination was performed by ranging the temperature from 20°C to 90°C with measurements every 2°C.

## Code availability

The code used for this study is available in the following repository https://github.com/bene837/af_gradmcmc

## Acknowledgments

The authors are funded by the European Research Council (Starting grant — 716058), the Swiss National Science Foundation, the National Center of Competence in Research in Molecular Systems Engineering, the high-performance computing facility at EFPL - SCITAS, the support from the Swiss Data Science Center and the CSCS Swiss National Supercomputing Centre.

## Supplementary information

**Figure S1.**
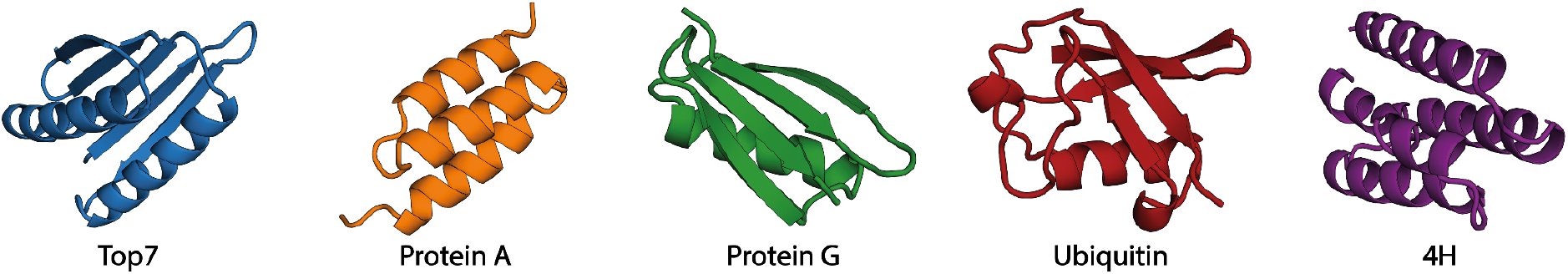
Target folds used for sequence design. Additional information about the structures can be found in supplementary table S4.

**Figure S2.**
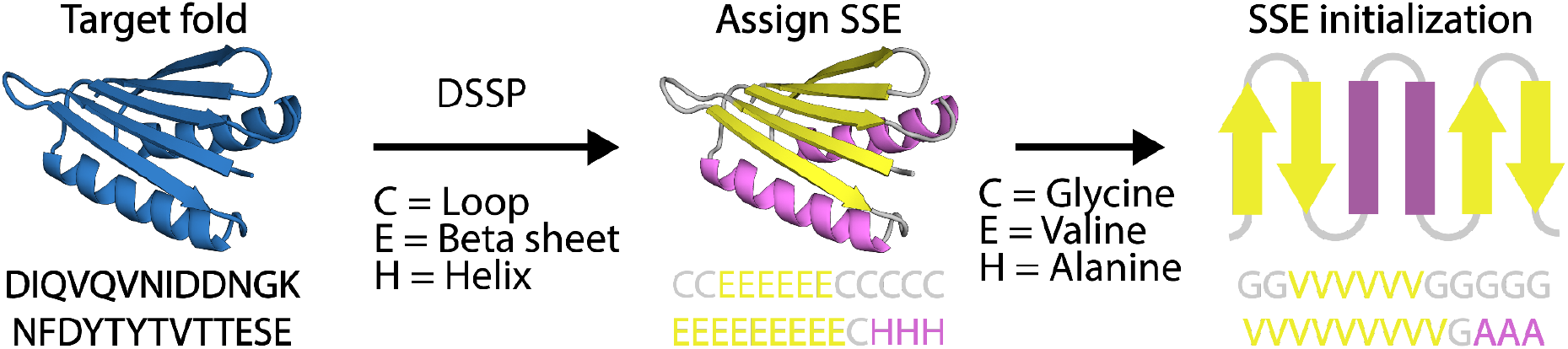
The Secondary Structural Element (SSE) initialization strategy. From a target fold the SSEs are assigned using Define Secondary Structure of Proteins (DSSP) (Kabsch & Sander, 1983). Next, amino acid identities are given to the residue positions given their SSE: helical residues get assigned alanines, beta-sheets valines and loops with glycines.

**Figure S3.**
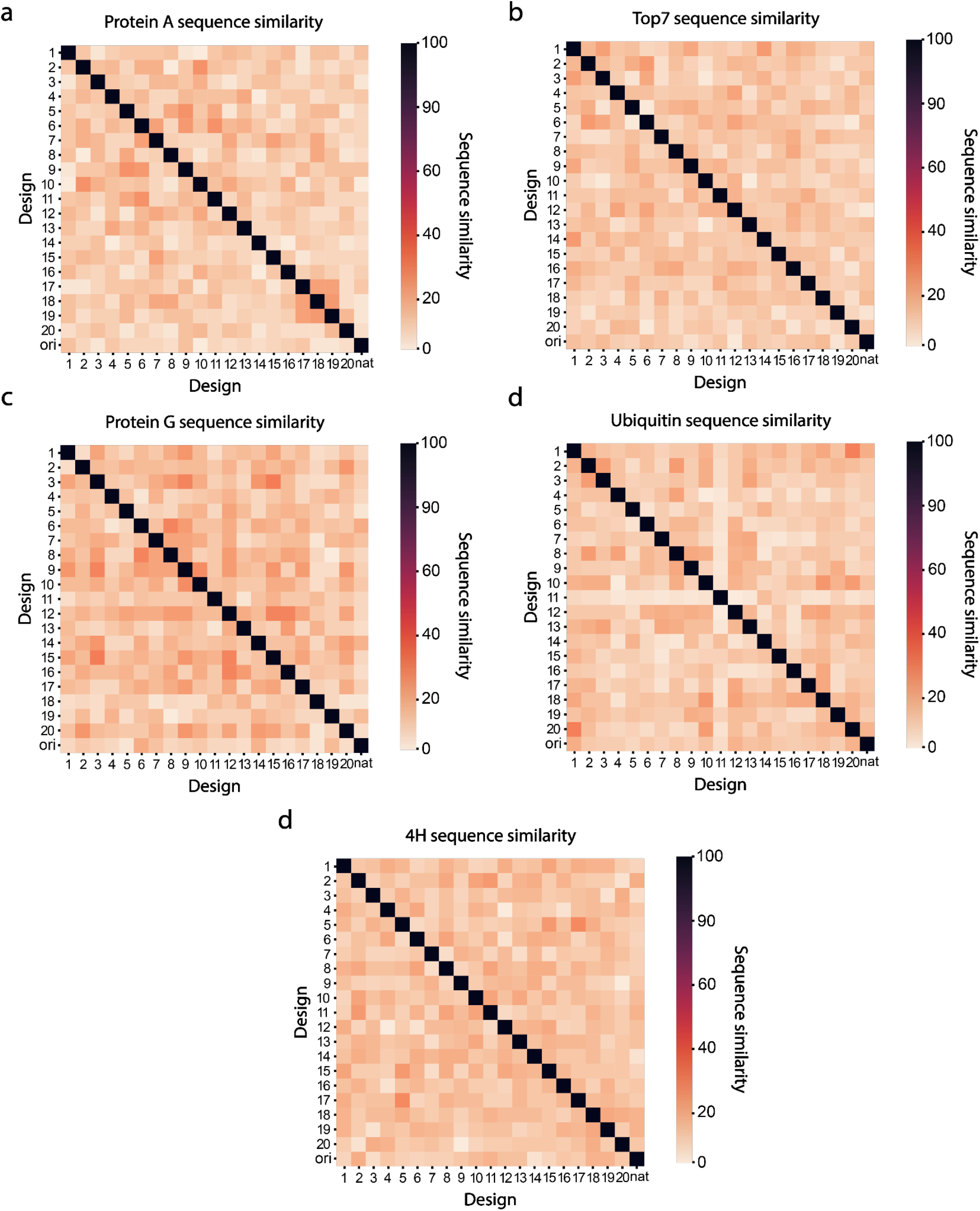
All vs all comparison of sequence similarity for 20 designs of each of the 5 folds. Designs are labeled 1-20 and ori is the native sequence of the design target.

**Figure S4.**
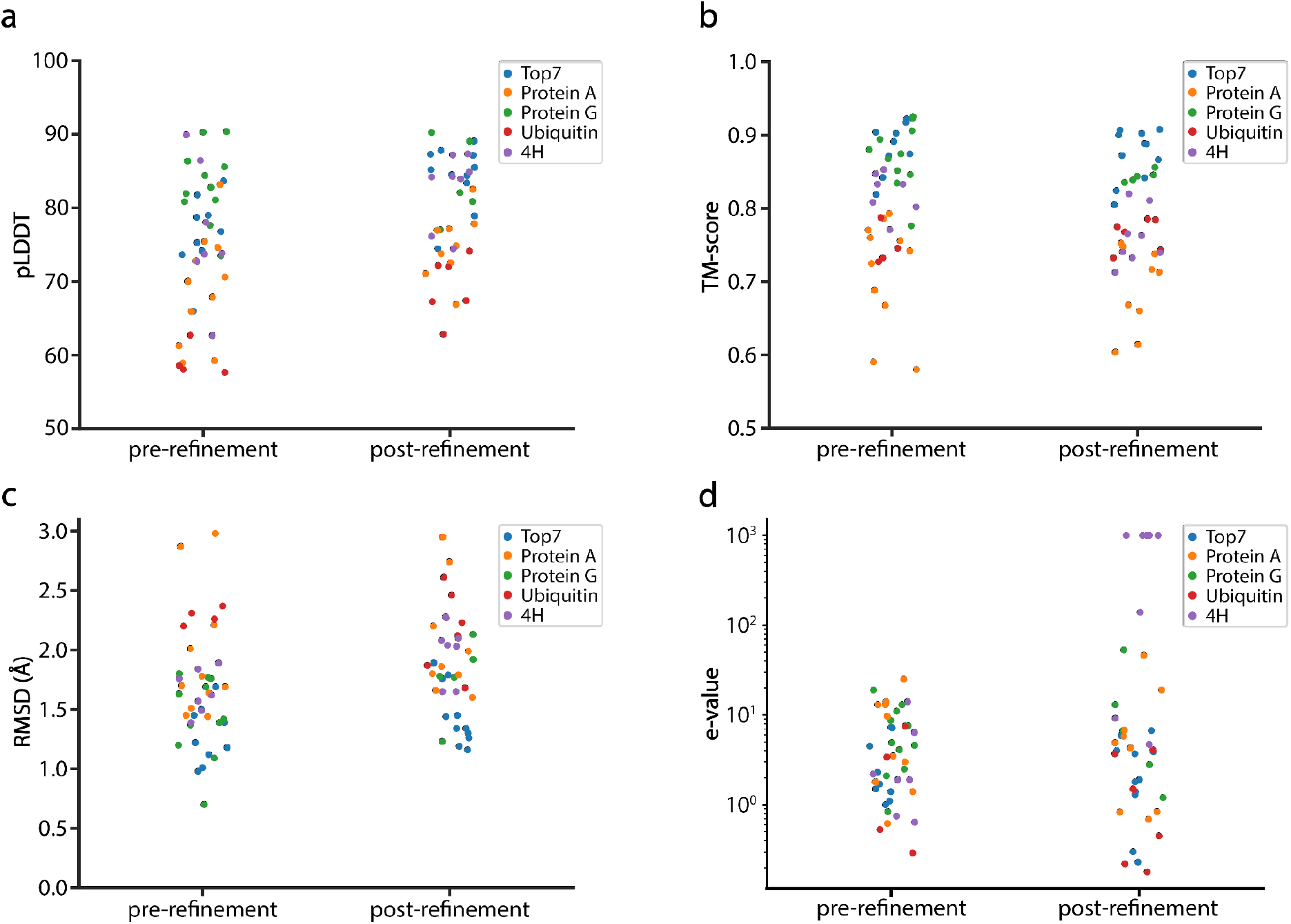
In silico metrics of the designs before and after Rosetta surface redesign. a) pLDDT scores before and after refinement as predicted by AF2. b) TM-scores of the predicted AF2 models vs. the target structure. c) RMSD of the predicted AF2 models vs. the target structure. d) E-values computed with BLAST before and after Rosetta surface redesign. E-values increase after sequence optimization to correct apolar aminoacids at the surface, moving further away from the natural protein sequence space. The e-values are cut off at a value of 1000.

**Figure S5.**
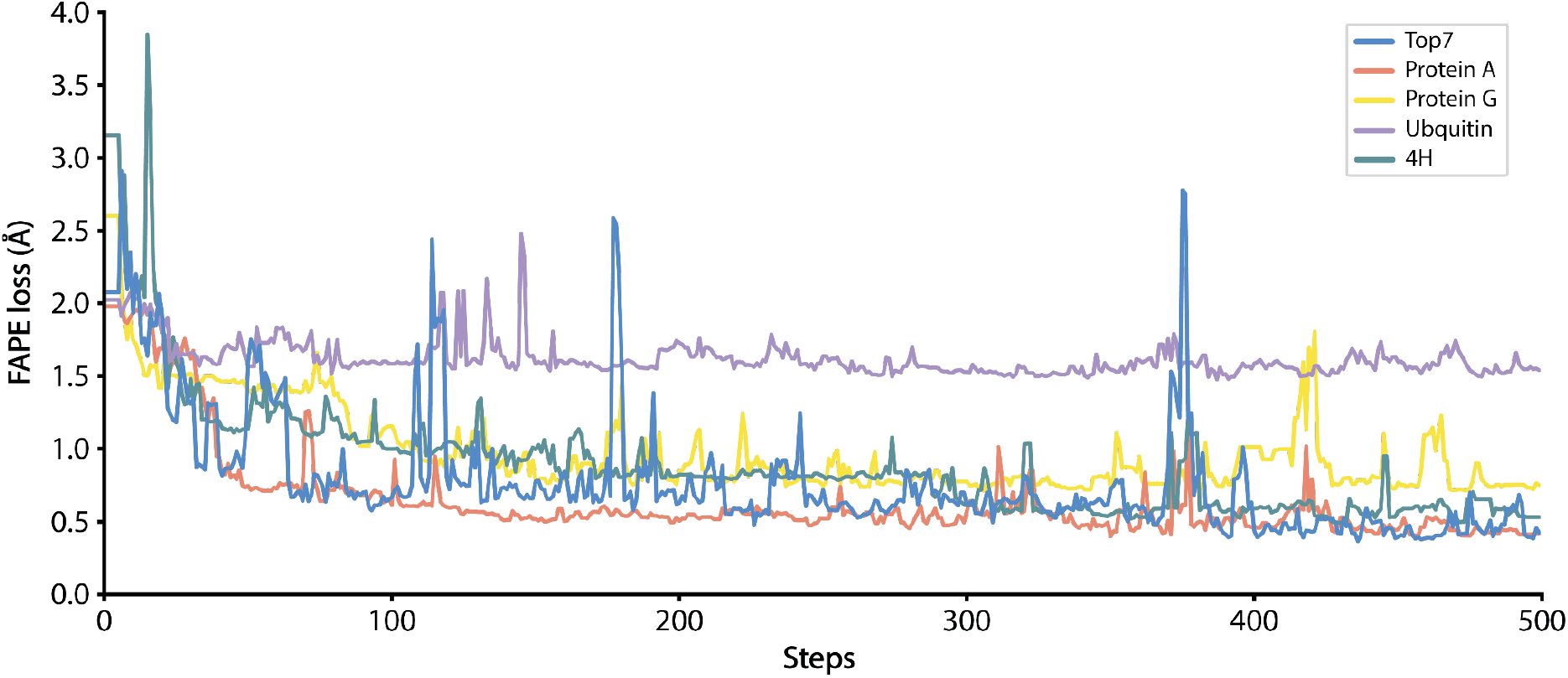
GD trajectories of the in vitro tested designs. For the designs tested in vitro one GD run was executed per target fold.

**Figure S6.**
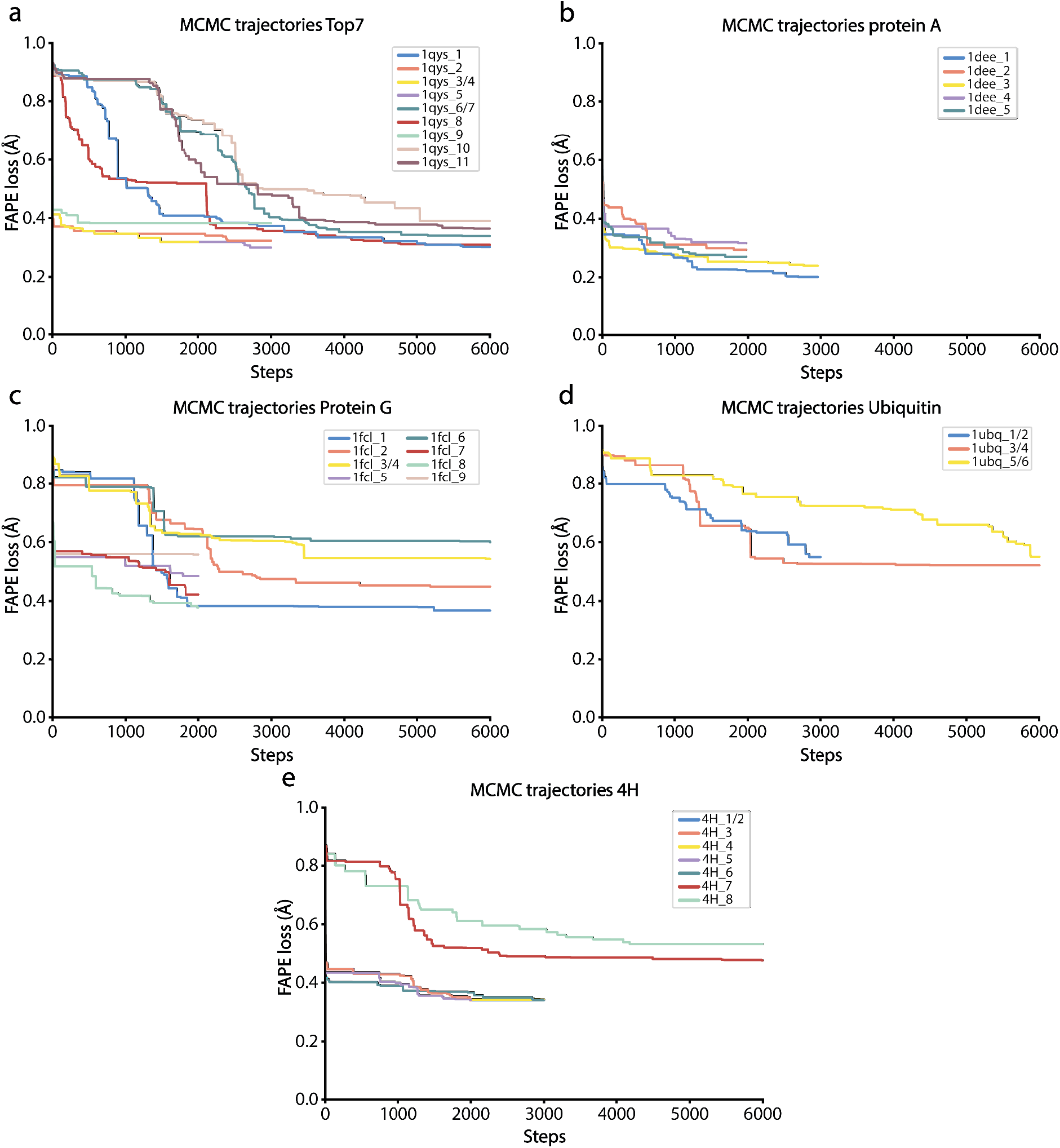
MCMC trajectories of the designs tested in vitro. MCMC trajectories were executed on the sequence designed by GD and on SSE-initialized sequences.

**Figure S7.**
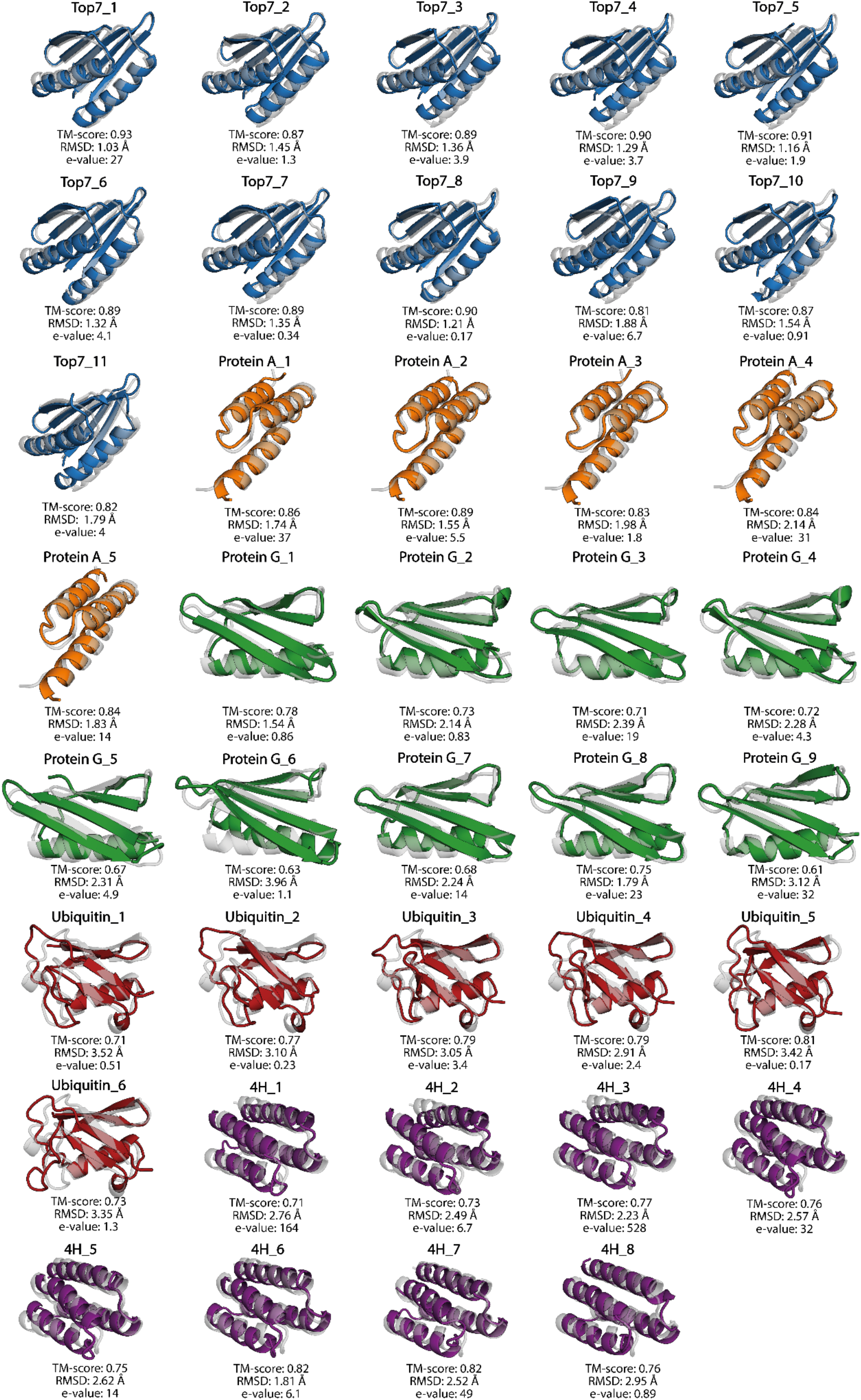
All AlphaFold2 models superimposed with the design target structure (gray). Top7 - blue, Protein A- orange, Protein G - green, Ubiquitin - red and 4H - purple.

**Figure S8.**
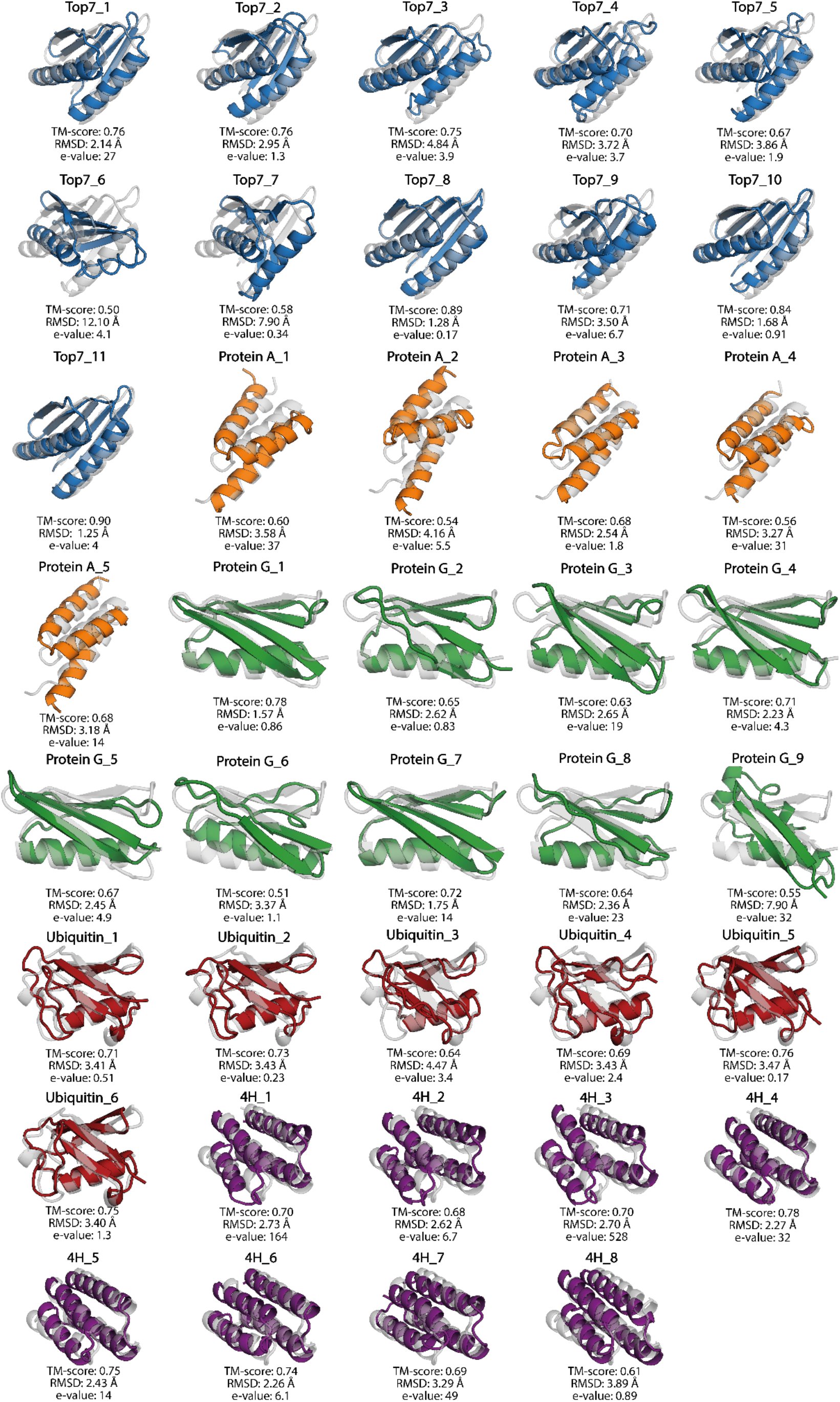
All RosettaFold models superimposed with the design target structure (gray). Top7 - blue, Protein A- orange, Protein G - green, Ubiquitin - red and 4H - purple.

**Figure S9.**
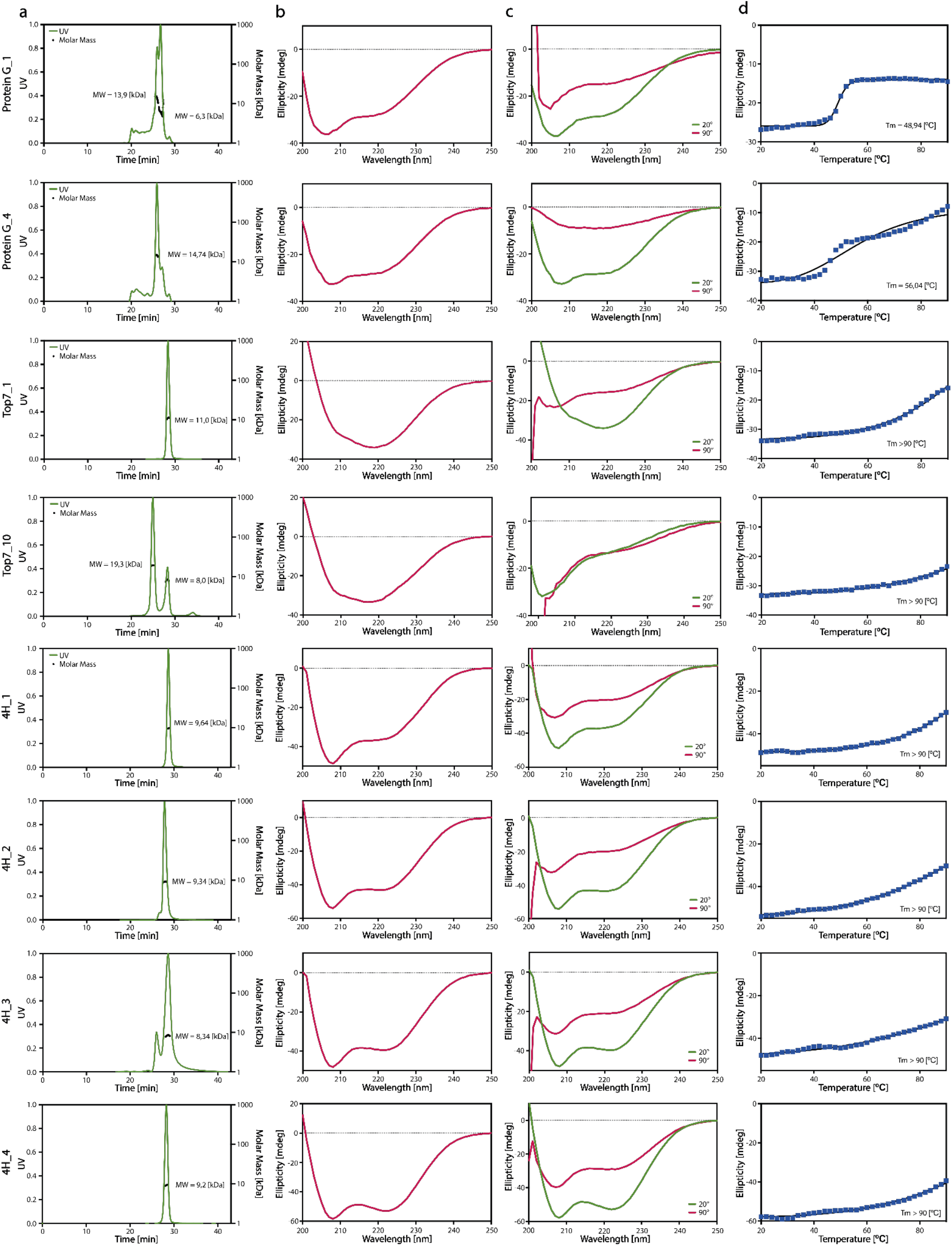
Biochemical characterization of the AF2 designs shown to be soluble and folded in solution. a) SEC-MALS measurements showing molecular weight determination. b) Circular dichroism spectra for assessment of secondary structure of the designs. c) Circular dichroism spectra at 20°C and 90°C. d) Temperature melting curves Thermal denaturation experiments to determine melting temperatures (T_m_).

**Figure S10.**
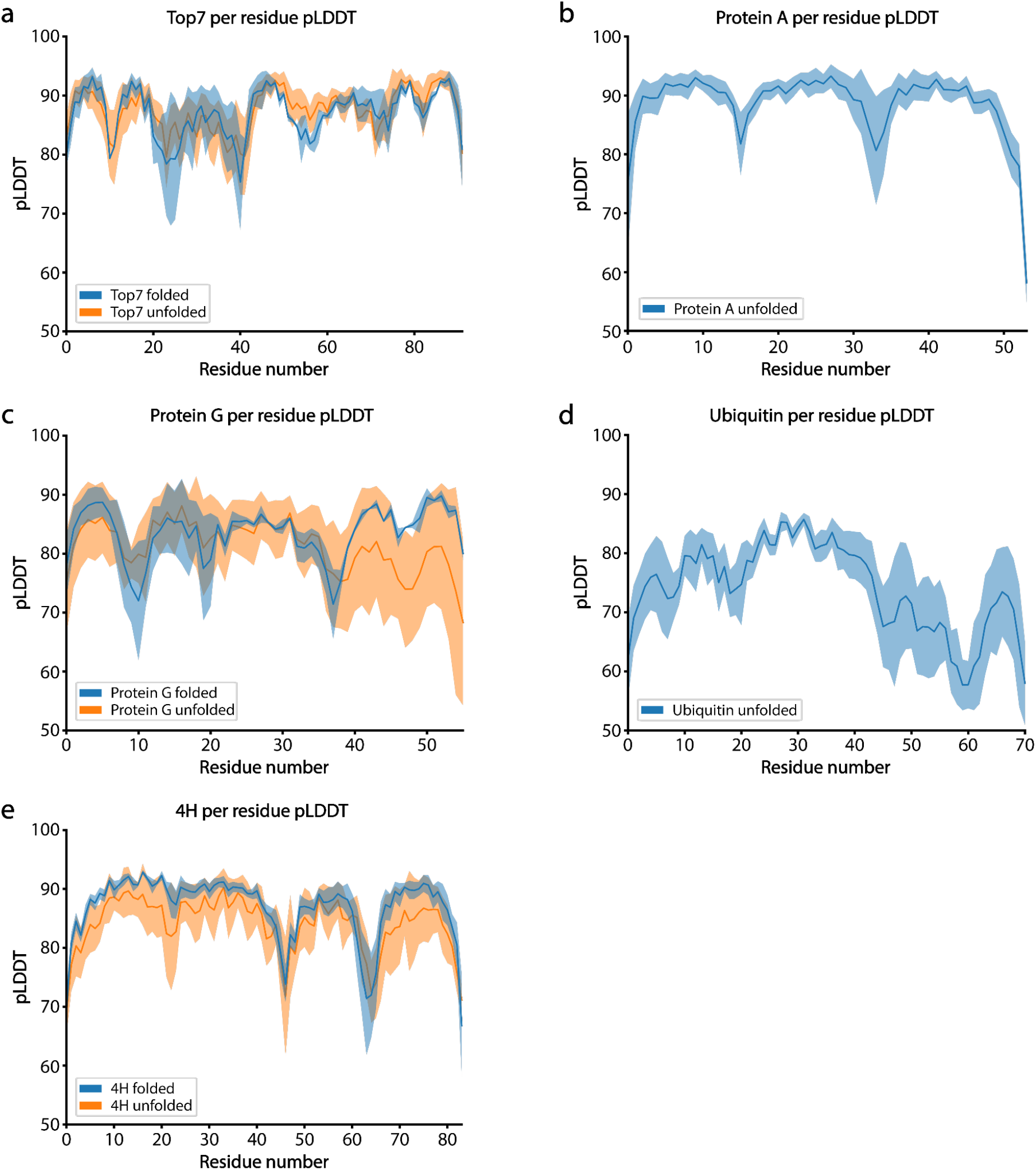
Confidence scores from AF2 per residue comparing folded vs unfolded designs. pLDDT scores from AF2 shown for both folded (blue) and unfolded (orange) designs tested in vitro.

**Figure S11.**
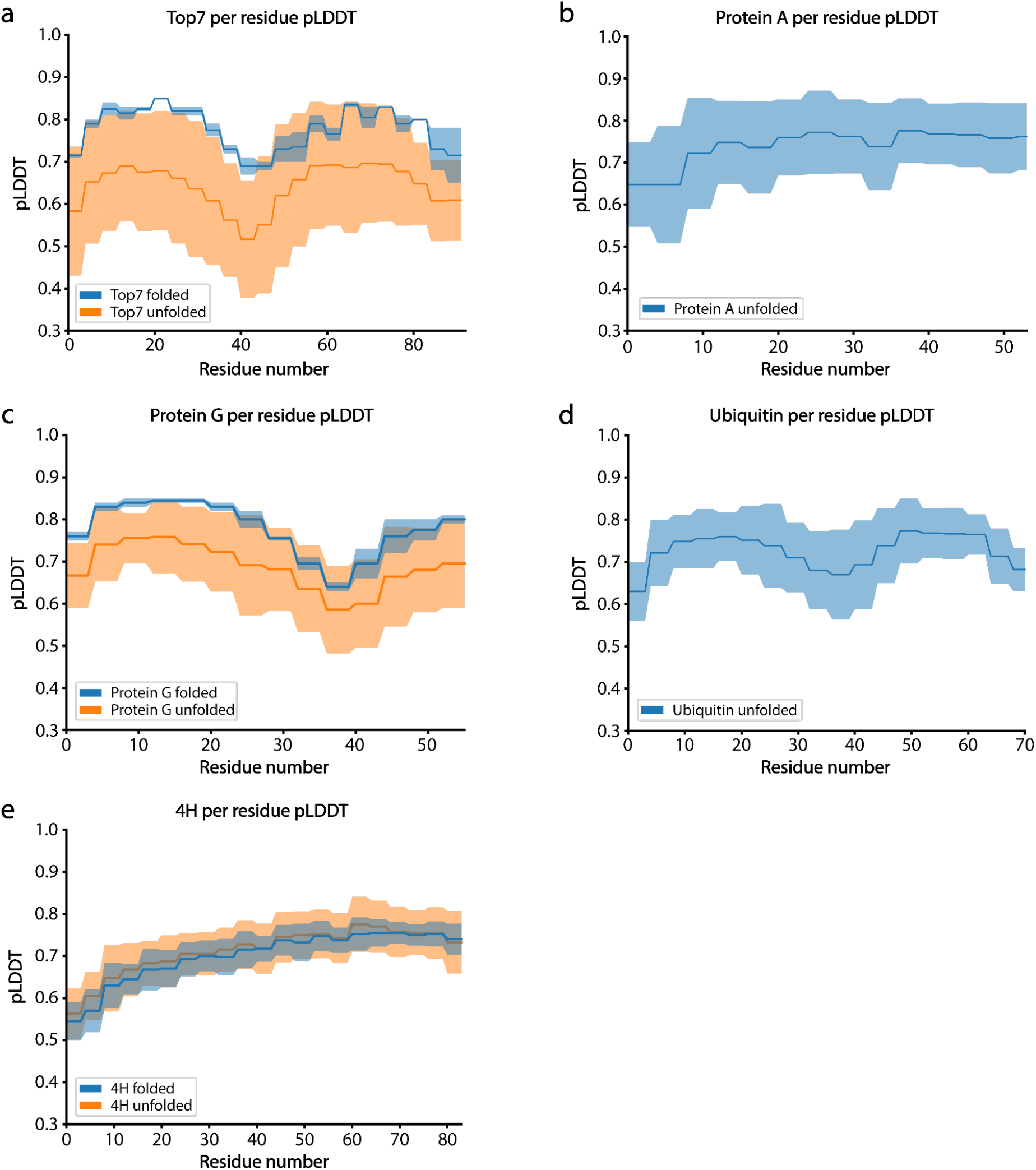
Confidence scores from RF per residue comparing folded vs unfolded designs. Per residue pLDDT scores from (RF) shown for both folded (blue) and unfolded (orange) designs tested of the in vitro validated designs.

**Table S1.**
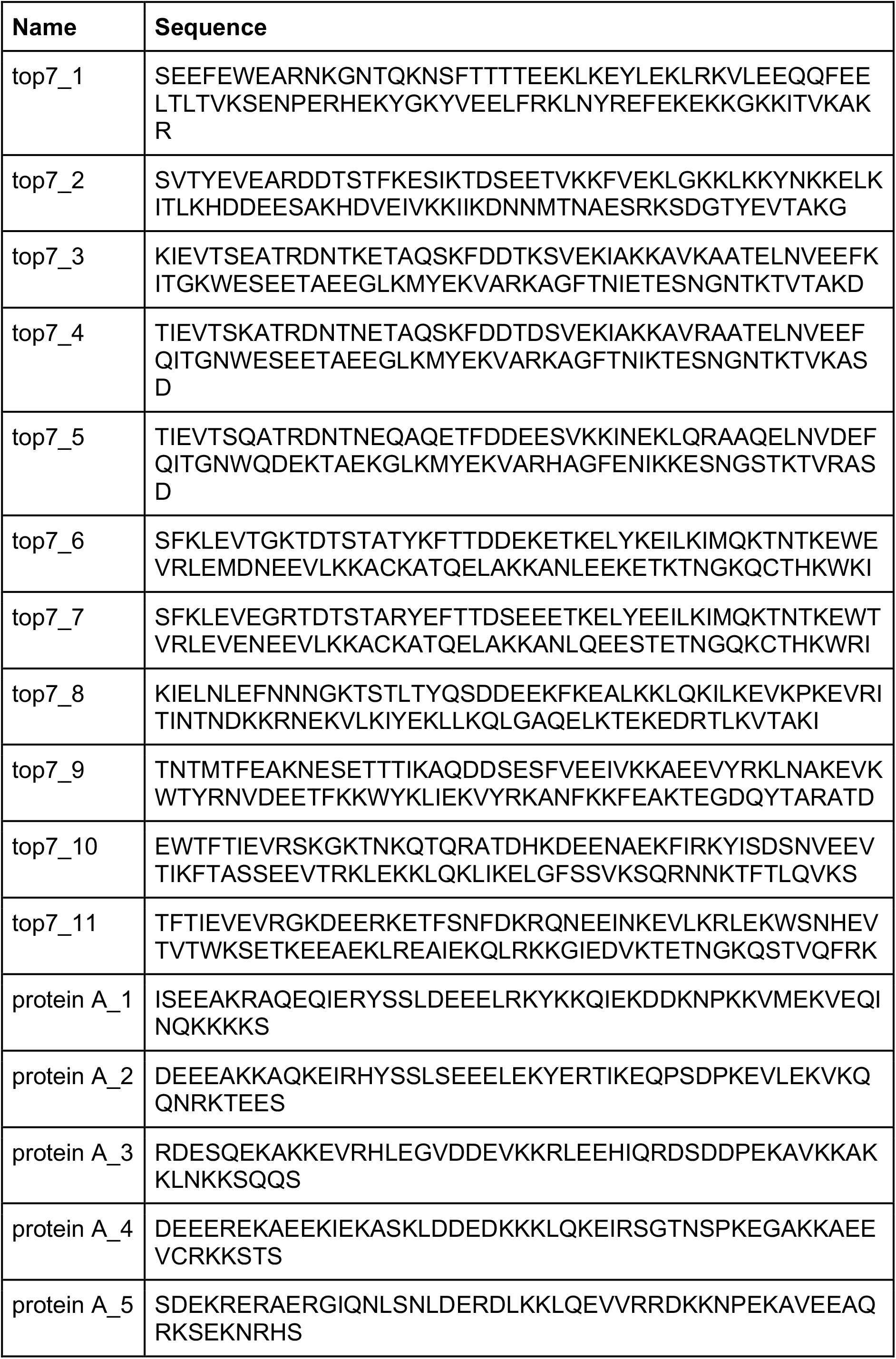
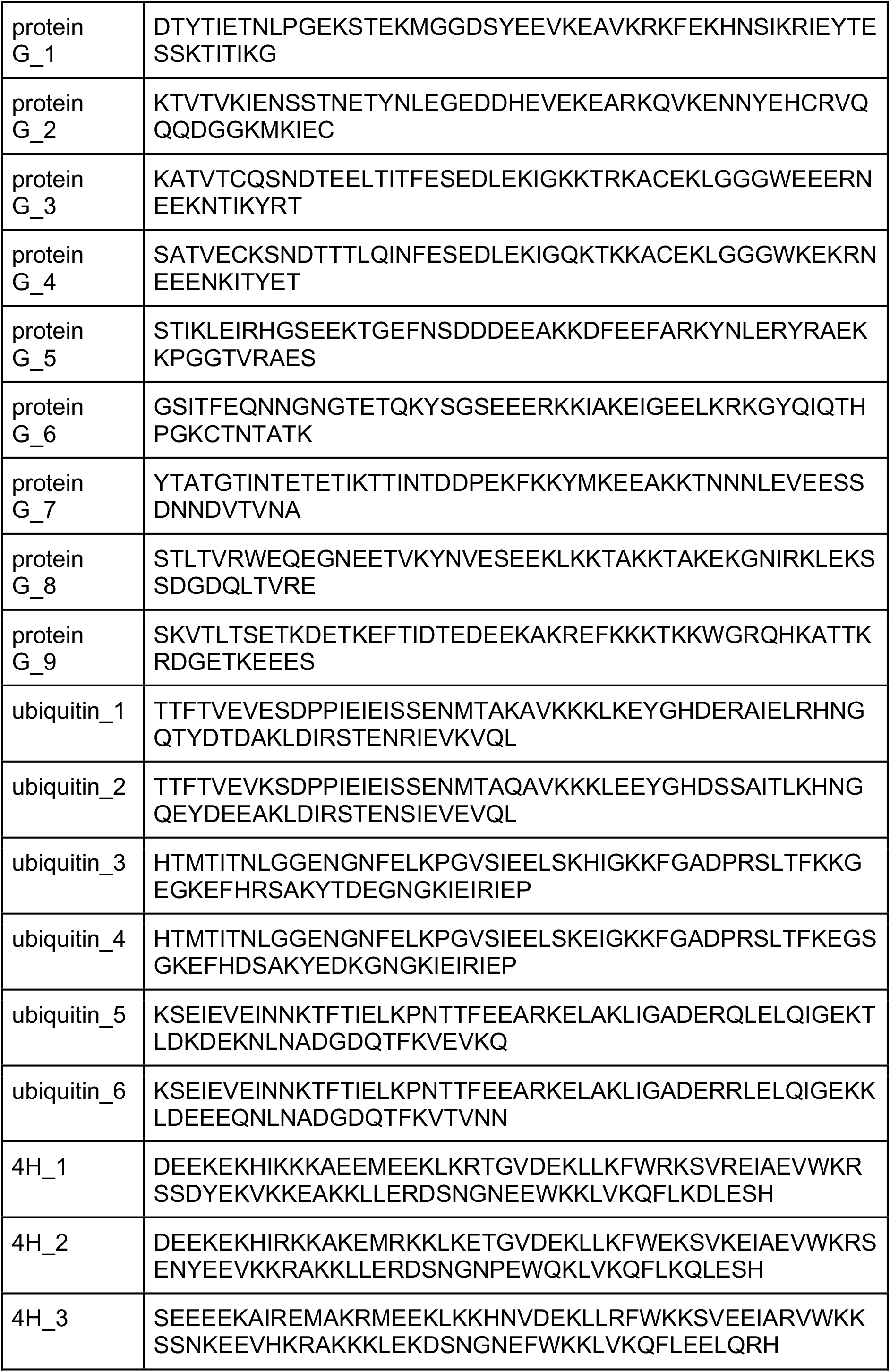
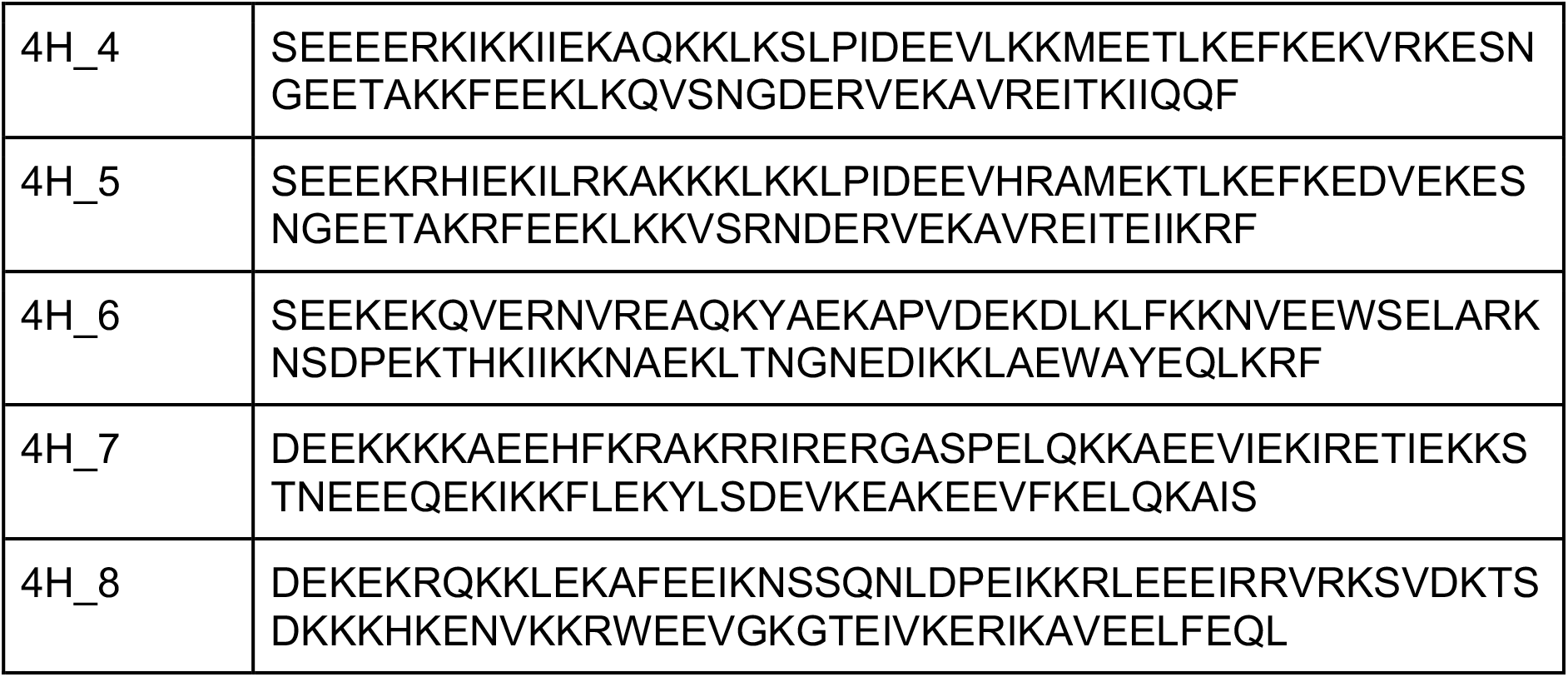
Sequences of the designs used for in vitro validation.

**Table S2.** AF2 and RF structural and confidence metrics of the designs used for in vitro validation.

| Name | pLDDT<br>(AF) | pLDDT<br>(RF) | TM-Score<br>(AF) | TM-Score<br>(RF) | RMSD<br>(AF) | RMSD<br>(RF) |
| --- | --- | --- | --- | --- | --- | --- |
| top7_1 | 88.1 | 0.77 | 0.93 | 0.76 | 1.03 | 2.14 |
| top7_2 | 83.4 | 0.72 | 0.87 | 0.76 | 1.45 | 2.95 |
| top7_3 | 87.8 | 0.59 | 0.89 | 0.75 | 1.36 | 4.84 |
| top7_4 | 78.9 | 0.67 | 0.90 | 0.70 | 1.29 | 3.72 |
| top7_5 | 74.5 | 0.64 | 0.91 | 0.67 | 1.16 | 3.86 |
| top7_6 | 85.1 | 0.60 | 0.89 | 0.50 | 1.32 | 12.10 |
| top7_7 | 86.5 | 0.60 | 0.89 | 0.58 | 1.35 | 7.90 |
| top7_8 | 85.7 | 0.82 | 0.90 | 0.89 | 1.21 | 1.28 |
| top7_9 | 87.3 | 0.69 | 0.81 | 0.71 | 1.88 | 3.50 |
| top7_10 | 84.0 | 0.76 | 0.87 | 0.84 | 1.54 | 1.68 |
| top7_11 | 84.5 | 0.79 | 0.82 | 0.90 | 1.79 | 1.25 |
| protein A_1 | 85.91 | 0.65 | 0.86 | 0.60 | 1.738 | 3.585 |
| protein A_2 | 76.77 | 0.67 | 0.89 | 0.54 | 1.55 | 4.159 |
| protein A_3 | 82.09 | 0.86 | 0.83 | 0.68 | 1.983 | 2.54 |
| protein A_4 | 90.67 | 0.78 | 0.84 | 0.56 | 2.138 | 3.267 |
| protein A_5 | 84.02 | 0.76 | 0.84 | 0.68 | 1.83 | 3.178 |
| protein G_1 | 75.68 | 0.76 | 0.78389 | 0.78 | 1.545 | 1.567 |
| protein G_2 | 77.15 | 0.77 | 0.73774 | 0.65 | 2.138 | 2.617 |
| protein G_3 | 77.83 | 0.74 | 0.7134 | 0.63 | 2.391 | 2.646 |
| protein G_4 | 71.07 | 0.77 | 0.71612 | 0.71 | 2.276 | 2.228 |
| protein G_5 | 66.88 | 0.71 | 0.66845 | 0.67 | 2.307 | 2.447 |
| protein G_6 | 85.39 | 0.56 | 0.62735 | 0.51 | 3.962 | 4.371 |
| protein G_7 | 73.38 | 0.74 | 0.68498 | 0.72 | 2.241 | 1.75 |
| protein G_8 | 78.87 | 0.71 | 0.75016 | 0.64 | 1.788 | 2.36 |
| protein G_9 | 76.52 | 0.43 | 0.60878 | 0.55 | 3.121 | 7.896 |
| ubiquitin_1 | 65.16 | 0.77 | 0.71 | 0.71 | 3.517 | 3.411 |
| ubiquitin_2 | 68.87 | 0.78 | 0.77 | 0.73 | 3.1 | 3.435 |
| ubiquitin_3 | 64.69 | 0.60 | 0.79 | 0.64 | 3.046 | 4.47 |
| ubiquitin_4 | 65.69 | 0.61 | 0.79 | 0.69 | 2.911 | 3.435 |
| ubiquitin_5 | 74.77 | 0.71 | 0.81 | 0.76 | 3.424 | 3.475 |
| ubiquitin_6 | 66.33 | 0.71 | 0.73 | 0.75 | 3.354 | 3.405 |
| 4H_1 | 84.16 | 0.66 | 0.71 | 0.70 | 2.764 | 2.73 |
| 4H_2 | 83.64 | 0.69 | 0.73 | 0.68 | 2.486 | 2.619 |
| 4H_3 | 86.89 | 0.68 | 0.77 | 0.70 | 2.226 | 2.697 |
| 4H_4 | 87.38 | 0.76 | 0.76 | 0.78 | 2.568 | 2.268 |
| 4H_5 | 84.77 | 0.77 | 0.75 | 0.75 | 2.619 | 2.432 |
| 4H_6 | 84.89 | 0.78 | 0.82 | 0.74 | 1.812 | 2.264 |
| 4H_7 | 76.88 | 0.62 | 0.82 | 0.69 | 2.524 | 3.292 |
| 4H_8 | 77.41 | 0.67 | 0.76 | 0.61 | 2.954 | 3.887 |

**Table S3.**
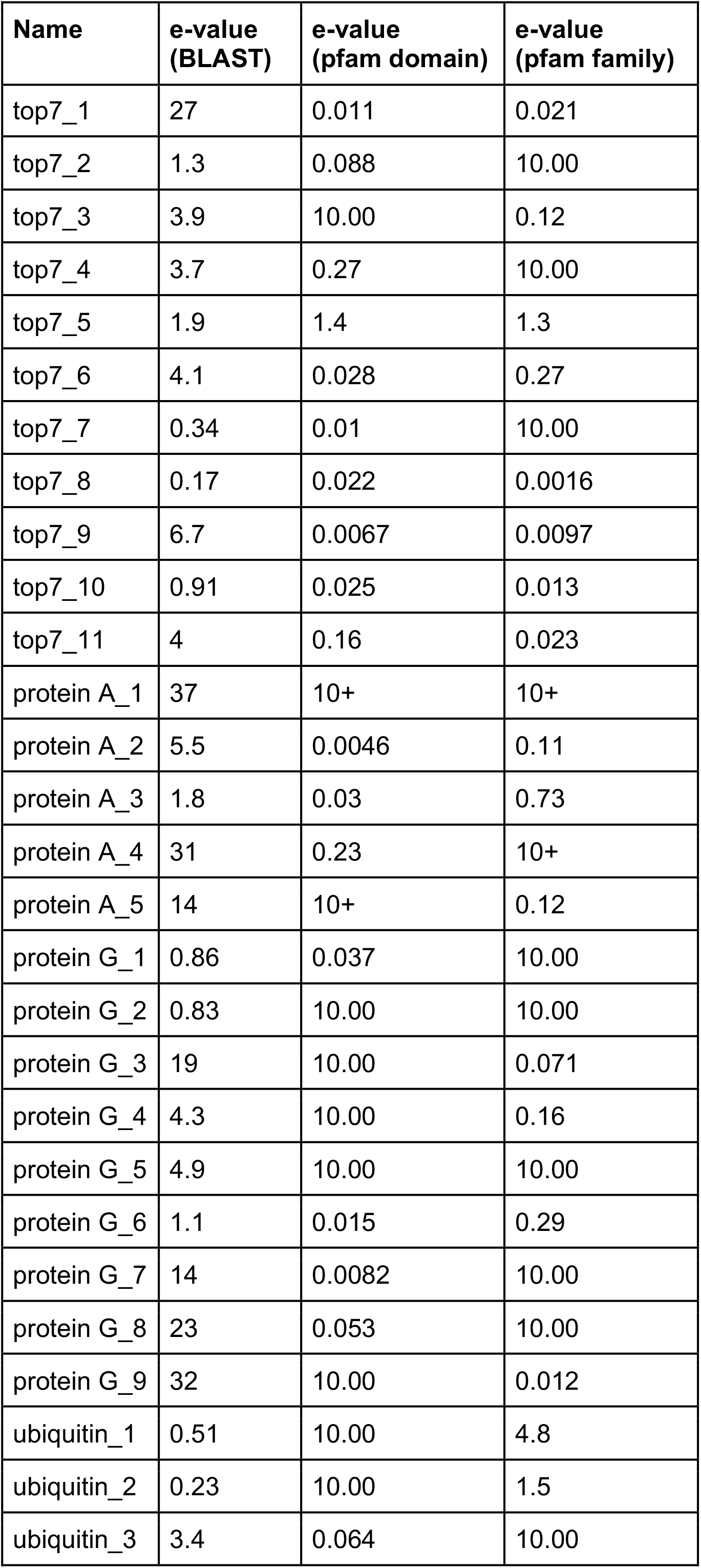
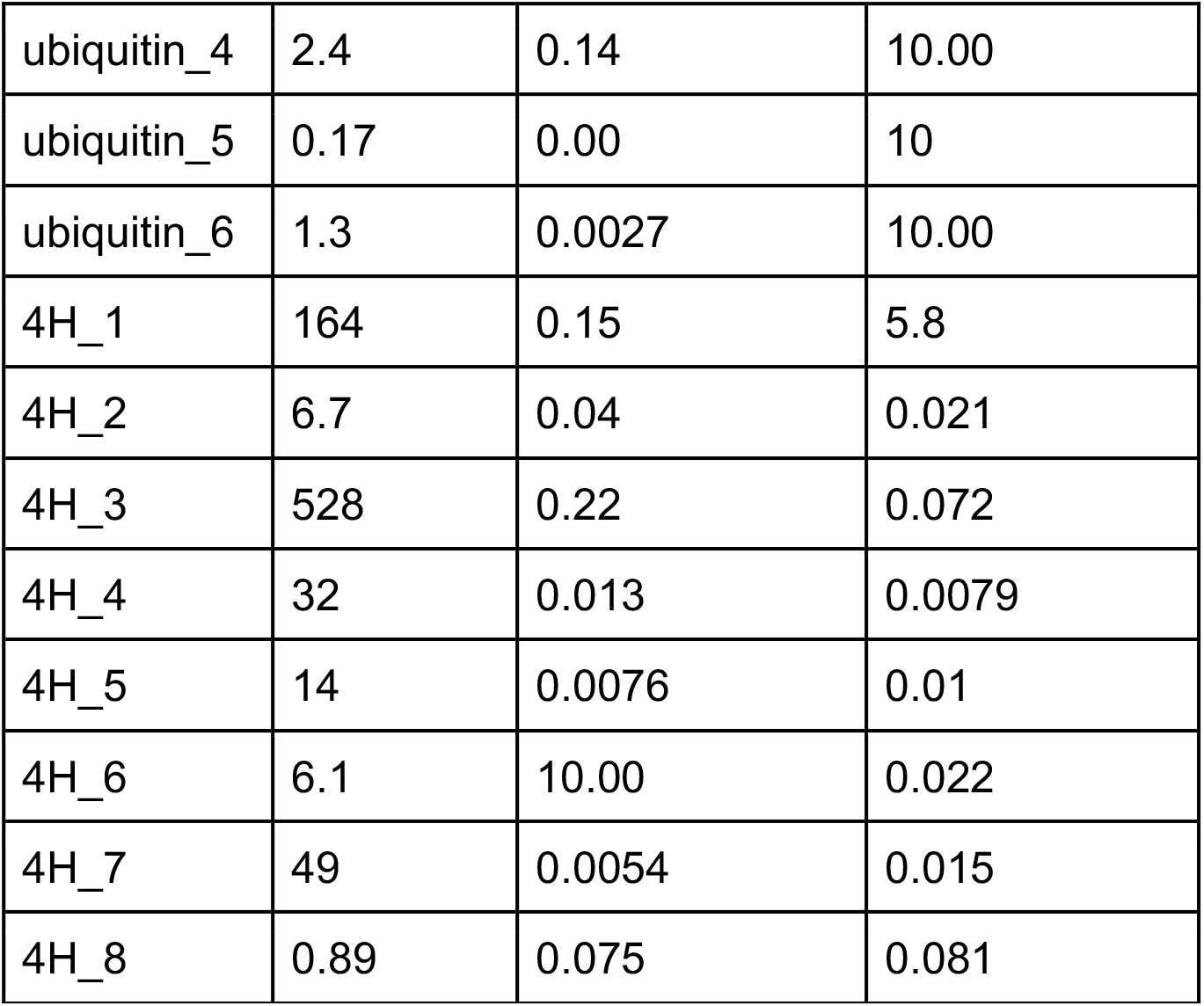
BLAST and pfam e-values of the designs used for in vitro validation. BLAST e-values have a maximum value of 1000 and pfam e-values of 10.

**Table S4.**
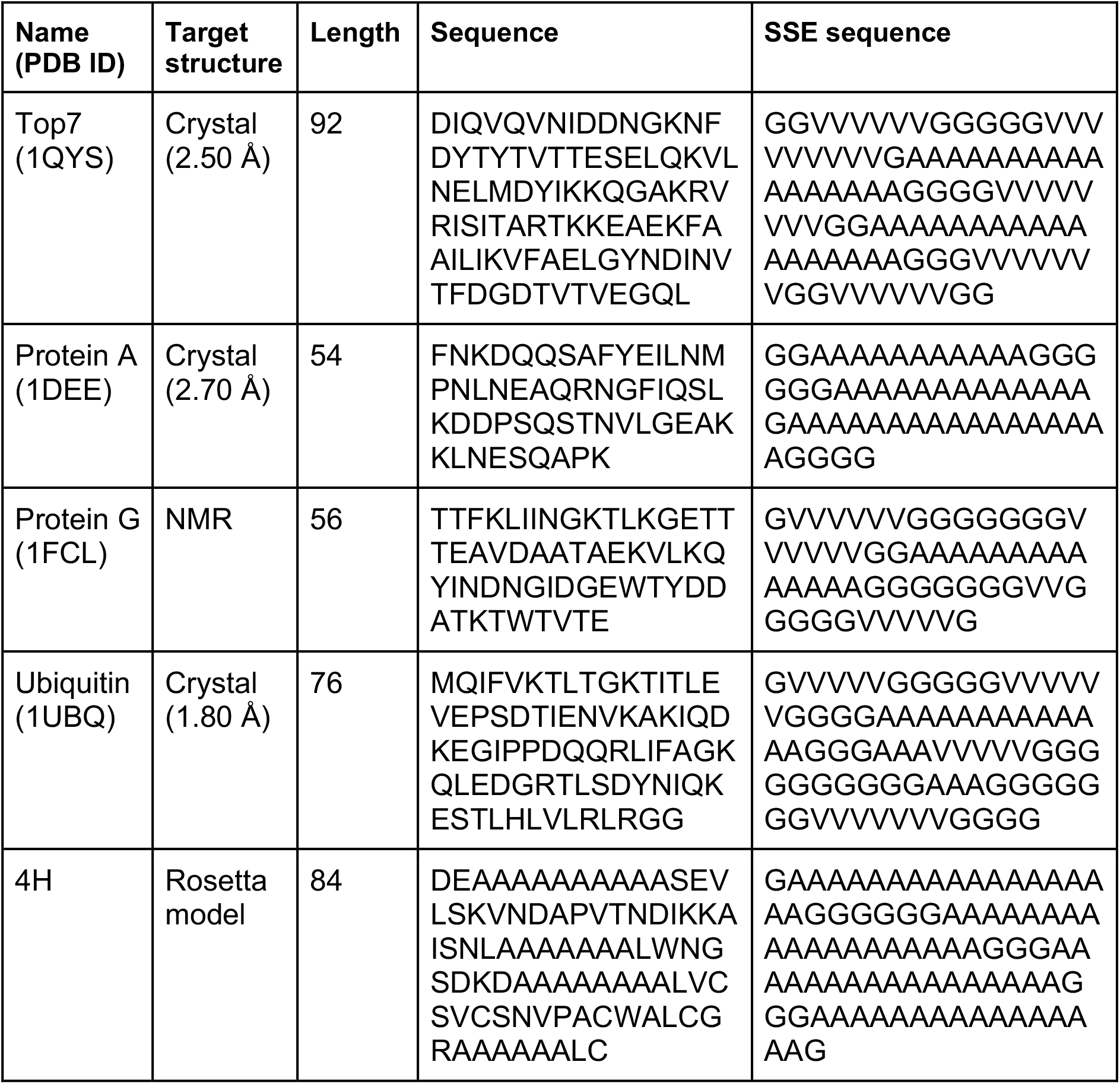
Information about the design targets. The sequence is obtained from the pdb file and SSEs are predicted using psipred and the SSE sequence is generated by placing alanines in helix positions, valines in beta sheet positions and glycine in loop positions.

